# Mifepristone alone and in combination with scAAV9-*SMN1* gene therapy improves disease phenotypes in *Smn^2B/-^* spinal muscular atrophy mice

**DOI:** 10.1101/2025.02.17.638672

**Authors:** Emma R Sutton, Eve McCallion, Joseph M Hoolachan, Özge Cetin, Paloma Pacheco-Torres, Sihame Bouhmidi, Lauren Churchill, Taylor Scaife, Helena Chaytow, Yu-Ting Huang, Stephanie Duguez, Bernard L Schneider, Thomas H. Gillingwater, Maria Dimitriadi, Melissa Bowerman

## Abstract

Spinal muscular atrophy (SMA) is a neuromuscular disease caused by deletions or mutations in the *survival motor neuron 1* (*SMN1*) gene. SMA is characterised by alpha motor neuron loss in the spinal cord and subsequent muscle atrophy. There are currently three approved SMN-directed therapies for SMA patients. While these therapies have transformed what was once a life-limiting condition into one that can be managed and even improved, they are unfortunately not cures, highlighting the need for additional supporting second-generation therapies. These should not only target the neuromuscular system but also peripheral and metabolic perturbations that are present in both SMA models and patients. *Krüppel-like factor 15* (*Klf15*) is a transcription factor that maintains metabolic homeostasis and is involved in the glucocorticoid-glucocorticoid receptor (GR) signalling pathway, in several peripheral and metabolic tissues in SMA mice. Here, we used murine and human cellular models as well as SMA mice and *Caenorhabditis Elegans (C. elegans)* to assess the therapeutic potential of reducing *Klf15* activity with mifepristone, a glucocorticoid antagonist, combined with SMN-targeted gene therapy. We report that mifepristone reduces *Klf15* expression across several *in vitro* models, ameliorates neuromuscular pathology in SMA *smn-1(ok355) C. elegans* and improves survival of SMA *Smn^2B/-^* mice. Furthermore, we show that combining mifepristone with an approved SMN-directed gene therapy (scAAV9-*SMN1*) results in improved tissue- and sex-specific responses to treatment. Our study demonstrates that a multi-tissue targeting SMN-independent drug, alone and in combination with an approved SMN-dependent therapy, has the potential to improve SMA disease pathology.

## INTRODUCTION

Spinal muscular atrophy (SMA) is a neuromuscular disorder characterised by the loss of alpha motor neurons in the anterior horn of the spinal cord and subsequent muscle atrophy ^1^. In addition to defects within the neuromuscular system; emerging studies have reported peripheral pathologies and metabolic perturbations in both human patients and mouse models ^2–4^. SMA is caused by a significant depletion but not complete loss of the survival motor neuron (SMN) protein ^5,6^. This is due to loss-of-function deletions and/or mutations in the *survival motor neuron 1* (*SMN1*) gene that are partially compensated by the presence of a second gene, the *survival motor neuron 2* (*SMN2*) gene, that is capable of producing ∼10% of fully functional SMN protein ^1,7^.

Ground-breaking and approved *SMN1* and *SMN2*-directed therapies for SMA (Spinraza, Zolgensma, Risdiplam) provide sustained improvement in motor function and increase lifespan of many patients, but these therapeutics are currently not a cure ^8^. As a result, the field is now looking beyond SMN and the neuromuscular system for additional contributors to pathology that may be targeted to provide additional therapeutic benefits ^9^. Peripheral pathologies in tissues such as the liver, heart, pancreas and skeletal muscle have been repeatedly reported in SMA mouse models and patients ^4,10–12^. Interestingly, these tissues play an important role in maintaining systemic energy homeostasis and their intrinsic defects in SMA could have significant consequences on whole-body metabolic homeostasis. Indeed, perturbations in fatty acid, amino acid, and glucose metabolism have been observed in SMA mouse models and patients ^3,10,13,14^. The fact that dietary supplementation improves lifespan of SMA mice ^15–17^ further supports the hypothesis that metabolic perturbations contribute to SMA pathology. More recently, we also demonstrated that providing an amino-based formula to children with SMA that had received an *SMN2*-directed treatment, significantly reduced their persisting gastrointestinal issues ^18^. Significant research also demonstrates additional peripheral comorbidities commonly reported in patients with SMA such as Metabolic Dysfunction-Associated Steatotic Liver Disease (MASLD; formally known as NAFLD) and type 1/ 2 diabetes ^4,19^. These studies highlight the importance and relevance of tackling peripheral and metabolic phenotypes in SMA which combines approved SMN-directed treatments and second-generation interventions ^20,21^.

An interesting and potential SMN-independent target is the transcription factor *Krüppel-like factor 15* (*Klf15*) ^3^. Transcriptional regulation is a main control mechanism of metabolic homeostasis and *Klf15*, a member of the zinc finger transcription factors, has an overarching influence on metabolic processes including those that are perturbed in SMA models and patients (fatty acid, amino acid and glucose metabolism) ^22–25^. The rhythmic expression of *Klf15* over a 24 hr period is modulated by the circadian secretion of glucocorticoids (GCs) and activity of the glucocorticoid receptor (GR) ^26,27^. *Klf15* then modulates several metabolic pathways, including the utilization of branched-chain amino acids (BCAAs) valine, leucine and isoleucine ^24^. We have previously shown an aberrant activity of the *Klf15*-GC-GR-BCAA pathway in serum and metabolic tissues from severe *Smn^-/-^;SMN2* and intermediate *Smn^2B/-^* SMA mice, whereby the levels of *Klf15* and GCs were elevated and BCAAs depleted ^3^. Importantly, modulation of the pathway by daily oral administration of BCAAs to *Smn^-/-^;SMN2* mice led to significant improvements in weight and survival ^3^.

The GC-GR-*Klf15*-BCAA pathway therefore contributes to SMA pathogenesis and could be a potential therapeutic target to alleviate peripheral and metabolic pathologies in SMA. As modulating BCAA metabolism, a downstream component of the GC-GR-*Klf15* signalling cascade, provided significant improvements in SMA mice, it is possible that targeting an upstream effector may lead to a more direct and specific modulation of the GC-*Klf15* pathway and thus, greater benefits. In this study, we therefore examined the therapeutic potential of mifepristone, a commercially available GR antagonist, for the treatment of SMA, alone and in combination with an SMN-dependent gene therapy. Furthermore, mifepristone appeared in a list of therapeutic candidates in our recently published study combining multi-omics and bioinformatics approaches to identify repurposed drugs for the management of muscle pathologies and SMA ^28^. Interestingly, mifepristone is currently being evaluated in clinical trials for several conditions including neurodegenerative and metabolic diseases such as cancer (ClinicalTrials.gov ID NCT03225547), Cushing’s syndrome (ClincalTrials.gov ID NCT00569582), Type 2 diabetes (ClinicalTrials.gov ID NCT05772169) and Alzheimer’s disease (ClinicalTrials.gov ID NCT00105105) ^29^.

In this study, we set out to determine the therapeutic potential of mifepristone in SMA. Our initial assessments found that mifepristone’s ability to reduce GC-induced *Klf15* expression was dependent upon cell type and differentiation state across representative immortalized cells lines for different metabolic tissue types (muscle, brown adipose tissue (BAT and liver). Importantly, we found that mifepristone treatment across SMA animals models including severe *Smn^-/-^;SMN2* SMA mice, milder *Smn^2B/-^* SMA mice and a severe*C. elegans* SMA model significantly improved disease phenotypes such as survival, muscle size and neuromuscular function. Finally, we assessed the potential synergistic activity of combining mifepristone with an *SMN1* gene-based therapy and observed tissue- and sex-specific effects. Overall, our study supports the relevance of using GC-antagonist drugs as secondary therapies alongside SMN-dependent treatments for targeting both neuromuscular and metabolic pathologies in SMA.

## MATERIALS & METHODS

### CELL CULTURE

#### Cell proliferation and differentiation

C2C12 and 3T3-L1 cells were maintained in growth media consisting of Dulbecco’s Modified Eagle’s Media (DMEM) (Gibco, Cat. #2041859). FL83B cells were cultured in Kaighn’s modification of ham’s F-12 media (ATCC, Cat. #30-2004). All cells were supplemented with 10% FBS (Gibco, Cat. #2025814K) and 1% penicillin/streptomycin (Gibco, Cat. #15140122). Cells were cultured at 37 °C with 5% CO_2_. C2C12 cells were differentiated in DMEM containing 1% FBS for 7 days. 3T3-L1 cells were differentiated in DMEM containing 1.0 μM dexamethasone, 0.5 mM methylisobutylxanthine (IBMX) (Sigma, Cat. #STBG0799V) and 1.0 μg/ml insulin (Sigma, Cat. #SLBW1822) for 48 hours followed by DMEM containing 10% FBS and 1.0 μg/ml insulin for 10 days, replenished every 2-3 days.

#### Cell drug treatment

Dexamethasone (Merck, D4902-100 mg) and mifepristone (SLS, M8046-100 mg) were diluted in ethanol. Cells were treated with 1 μM, 5 μM and 10 μM dexamethasone for either 4, 8 or 24 hours. Optimal concentration of dexamethasone was combined with mifepristone at 1 μM, 5 μM and 10 μM for varying time points dependent on preliminary experiments (4, 8 or 24 hours).

#### Lactate dehydrogenase (LDH)-Glo^TM^ cytotoxicity assay

Cell toxicity was determined using the Lactate dehydrogenase (LDH)-Glo^TM^ assay kit (Promega) as per manufacturer’s instructions. Briefly, cells were treated with mifepristone (10 μM) and vehicle for either 24 or 72 hours. Controls included no cell control, vehicle only cell control and maximum LDH release control. Max LDH control cells were exposed to 10% Triton X-100 for 15 minutes. For all cell lines, a 1:300 dilution was used for undifferentiated cells and a 1:100 dilution was used for differentiated cells. The assay reaction was performed in the dark, at room temperature in 96-well opaque plates using 50 μl LDH detection reagent (1:200 reductase to detection enzyme mix) and 50 μl sample media (1:1). Luminescence was recorded after 60 minutes on a GloMax Explorer (Promega).

#### Bromo-2-deoxyuridine (BrdU) assay

Cell proliferation was determined by using 5-Bromo-2-deoxyuridine (BrdU) colorimetric system (Merck) as per manufacturer’s instructions. Briefly, cells were exposed to mifepristone (10 μM) or vehicle (ethanol) for 8 hours prior to BrdU assay. The cells were labelled with BrdU (1:2000) for 16 hours. Cells were exposed to an anti-BrdU fluorescence-labelled antibody. Using a spectrophotometric plate reader (GloMax Explorer (Promega)), BrdU absorbance was measured at dual wavelengths of 450-600 nm.

### ANIMALS AND PROCEDURES

#### Study approval

The *Smn^2B/-^* mouse line was housed at the Keele University Biomedical Sciences Unit (BSU) and cared for according to Home Office Animal Scientific Procedures Act 1986 (ASPA) regulations (project license: P99AB3B95, personal license: IO376FCD7). All procedures were approved by the Keele University ethics committee (AWERB). Mouse models *Smn^2B/-^, Smn^2B/2B^* mice (obtained from Charles River) and *Smn^+/-^* mice (obtained from Jackson labs) were crossed to obtain *Smn^2B/-^* and *Smn^2B/+^* mice. *Smn^+/-^* mice were crossed with *Smn^+/-^* mice to obtain C57BL/6J wild-type mice. Genotyping was performed on ear clips by PCR. Treatment groups were assigned at random, and animals of both sexes were used in all experiments.

The FVB.cg-*SMN1*tm1HungTg(*SMN2*)2Hung/J mice were crossed with *Smn^+/-^* mice producing 50% SMA offspring (*Smn^-/-^;SMN2tg/0*) and 50% control carriers (*Smn^-/+^;SMN2tg/0*) ^30^. These ‘Taiwanese’ SMA mice were housed in the University of Edinburgh animal facilities, in a 14-hour/10 hour light/ dark cycle in individually ventilated cages. All procedures were conducted according to Home Office ASPA 1986 regulations (Project License: PP1567597; PILs: IAC4805FD and 7E4CB171).

#### Drug administration in Smn^2B/-^ mice

Mifepristone (RU486) (SLS, Cat. #M8046-100MG) was solubilised in 2ml 0.5% carboxymethylcellulose (CMC) and sonicated for 3 minutes at 37kHz. Mifepristone was administered by oral gavage at concentrations of 250 μg/g from post-natal day 8 (P8) or 500 μg/g starting from P5. The scAAV9-*SMN1* vector was produced by transient transfection of HEK293 cells adapted to suspension culture (HEKExpress^TM^, ExcellGene SA) and the AAV9 particles were purified from the cell pellet and supernatant using affinity chromatography (POROS^TM^ CaptureSelect^TM^ AAV9 affinity resin; ThermoFisher Scientific) ^31^. After concentration, the vector titer measured by dPCR was 1.4E13 VG/mL. The vector was administered by facial vein injections to post-natal P0 pups (1E11 VG/pup, 20μl volume/pup). Combinatorial treatment with scAAV9-*SMN1* was administered by facial vein injections to P0 pups (1E11 VG/pup, 20μl volume/pup) combined with 500 μg/g mifepristone by daily oral gavage starting at P5 until P21.

Phenotypic analysis of weight and righting reflex was conducted daily on all mice. Survival analyses were conducted on litters until defined humane endpoints were reached. For all experiments litters, containing males and females, were randomly assigned treatment. Triceps, tibialis anterior (TA), liver and BAT were harvested for molecular and histological analysis from *Smn^2B/-^* mice and *Smn^2B/+^* healthy littermates.

#### Drug administration in Taiwanese Smn^-/-^;SMN2 mice

Mifepristone was prepared as above. Phenotypic analyses were conducted daily on *Smn^-/-^ ;SMN2* mice. Litters undergoing daily 500 μg/g mifepristone treatment versus vehicle control were administered the drug on P3 by oral gavage up until animals reached their humane endpoint.

Litters undergoing daily 500 μg/g mifepristone treatment were administered the drug on P3 by oral gavage up until animals reached their humane endpoint. Procedures were performed in a laminar flow hood in the animal facility by Helena Chaytow.

#### C. elegans SMA model

The LM99 *smn-1*(ok355)/*hT2* strain, segregates into homozygotes *smn-1*(ok355), lethal homozygotes *hT2/hT2*, and heterozygotes *smn-1(ok355)/hT2*. Homozygotes *smn-1*(ok355) resemble a severe SMA model. Heterozygotes smn-1/hT2 were used as controls. These animals were maintained at 20◦C on Nematode Growth Medium (NGM) plates seeded with Escherichia coli OP50 bacteria ^32^.

Mifepristone (RU486) (SLS, Cat. #M8046-100MG) was dissolved in DMSO and added to the NGM agar solution at concentrations of 0, 1, 15 and 30 μM. Mifepristone was administered by to *C. elegans* by raising the animals on plates containing the vehicle or drug.

Neuromuscular assays were performed on *C. elegans* that were 3 days old. The pharyngeal pumping assay was performed as previously described ^33^. Notably, an Axio Cam ICc5 camera on a Discovery V8 SteREO microscope was used for both movement assays. The pharyngeal pumping assay was filmed using 150X objective at 175 frames/10 seconds. A grinder movement in any axis was defined as a pumping event. Pumps were manually counted using Zen Pro software v2.3. Locomotion assays were filmed using a 63X objective at 15 frames/second. Mobility forward time, for 5 minutes, was quantified using WormLab 1.1 software (MBF Bioscience).

#### Laminin staining of skeletal muscle

TA muscles were fixed in 4% paraformaldehyde (PFA) overnight. Tissues were mounted in cryomoulds and quickly frozen in liquid nitrogen. Tissues were sliced at 13 μM and stored at −20 °C. Briefly, sections were dipped in acetone for 5 minutes and left to air dry for 30 minutes and incubated for 2 hours in blocking buffer (0.3% triton X, 20% FBS, 20% BSA in PBS). Samples were incubated overnight at 4°C with a rat anti-laminin antibody (1:1000, L0663, Sigma Aldrich) in blocking buffer. Followed by goat-anti-rat IgG 488 secondary antibody (1:250, AlexaFluor488, ThermoFisher scientific) for one hour. Tissues were mounted in media containing DAPI (SLS, F6057-20ML). Images were taken with a fluorescence microscope. TA muscle fibre area was measured on at least100 fibres from 3-5 sections per animals using Fiji. Images were assigned a random ID number by another experimenter and true IDs were only revealed once quantification was finalised.

#### Primary type III SMA human deltoid myoblasts

Dr Stephanie Duguez (Ulster university) generously provided cell pellets and/or RNA samples from SMA Type III primary human myoblasts and age-matched healthy controls obtained from deltoid muscle biopsies.

#### qPCR

RNA was extracted from cells using the ISOLATE II RNA Mini Kit (Bioline), following manufacturer’s instructions. Tissues (triceps, BAT and liver) were homogenised in RLT buffer using 7 mm stainless steel balls (Qiagen) and the Tissue Lyser LT (Qiagen) set at 50 oscillations for 2 minutes. Extraction of RNA from skeletal muscle was conducted using the RNeasy Fibrous tissue kit (Qiagen). The ISOLATE II RNA mini kit (Qiagen) was used to extract RNA from liver, while BAT RNA was extracted using the RNeasy lipid tissue mini kit (Qiagen), as per manufacturer’s instructions. A Nanodrop 1000 spectrophotometer (ThermoScientific) was used to measure the RNA concentrations (ug/ul) of samples alongside a blank control sample using RNase-free water. cDNA was prepared using cDNA synthesis mix (4 μl) and 20x RTase (1 μl). cDNA was then produced by reverse transcription using a ^3^Prime Thermocycler (Techne). qPCR was performed on StepOnePlus™ Real-time PCR system (ThermoFisher Scientific). *PolJ* was used as a housekeeping gene and relative gene expression was quantified using the Pfaffl method and primer efficiency was calculated using LinRegPCR V11.0 software. The list of mouse and human primers used can be found in Supplementary Table 1.

#### Statistical analysis

The most up to date GraphPad Prism software was used for data analysis and are presented as the mean ± standard error the mean. Appropriate statistical tests were used depending on data set, including unpaired t-test, one-way analysis of variance (ANOVA) and two-way ANOVA followed by post-hoc tests. Kaplan-Meier survival analysis was performed using the log-ran (Mantel-Cox) test. Statistical significance was found with P values less than 0.05 displayed as \**P*< 0.05, \*\**P*< 0.01, ***P< 0.001, \*\*\*\**P*< 0.0001. Outliers were identified using the GraphPad Grubb’s Test Individual data sets were all ran through the Grubbs’ test calculator, with the significance level set to alpha 0.05, to determine whether the most extreme value in the data set was a notable outlier compared to the other values. If this was the case, outliers were excluded from the analysis.

## RESULTS

### Mifepristone reduces *Klf15* expression in cellular models of metabolic tissues

We initially assessed mifepristone activity in immortalised cells that reflect metabolic tissues in which we had reported increased expression of *Klf15* ^3^: the C2C12 cell line for skeletal muscle, the 3T3-L1 cell line for brown adipose tissue (BAT) and the FL83B cell line for liver tissue^34–36^.

To model the increase in *Klf15* expression seen in tissues of SMA mice ^3^, we used a synthetic glucocorticoid, dexamethasone, to induce *Klf15* expression. To determine the optimal dosing regimen for maximal expression of *Klf15*, we treated cells with different concentrations of dexamethasone (1, 5 and 10 μM) across different time points (4, 8 and 24 hours) (Supplementary Figure 1A-C). We therefore determined that the optimal treatment regimens were: 10 μM for 24 hours for both differentiated C2C12 myotubes and FL83B hepatocytes as well as 10 μM for 4 hours for 3T3-L1 brown adipocytes (Supplementary Figure 1A-C).

For each cell line, we assessed the expression of both GR isoforms, whereby GRα is the main mediator of GCs while GRβ inhibits GRα and induces GC resistance ^37,38^. We then determined mifepristone’s ability to reduce *Klf15* expression by adding it (1, 5 and 10 μM) before or after the previously determined optimal dexamethasone treatment. In addition, we evaluated the effect of mifepristone on cell death and proliferation, using the lactate dehydrogenase (LDH)-Glo™ assay and a Bromo-2-deoxyuridine (BrdU) colorimetric assay, respectively.

In differentiated C2C12 myotubes (D7), we found a significant increase in the expression of both GRα and GRβ isoforms compared to proliferating myoblasts (D0) (Figure 1A), suggesting differential glucocorticoid sensitivities throughout muscle development. Next, we investigated the ability of mifepristone to reduce dexamethasone-induced *Klf15* expression in C2C12 myotubes. We found that mifepristone treatment alone in differentiated C2C12 myotubes significantly reduced *Klf15* levels at all three doses used when compared to untreated cells (Figure 1B). Interestingly, mifepristone, whether added before or after dexamethasone, did not reduce *Klf15* expression in C2C12 myotubes when compared to dexamethasone-treated cells (Figure 1C). Finally, the highest dose of mifepristone (10 μM) did not significantly impact cell death of C2C12 myotubes (Figure 1D) or proliferation of C2C12 myoblasts (Figure 1E) as compared to untreated cells, suggesting that although mifepristone treatment was safe, it did not reduce *Klf15* expression in GC-treated C2C12 myotubes.

**Figure 1.**
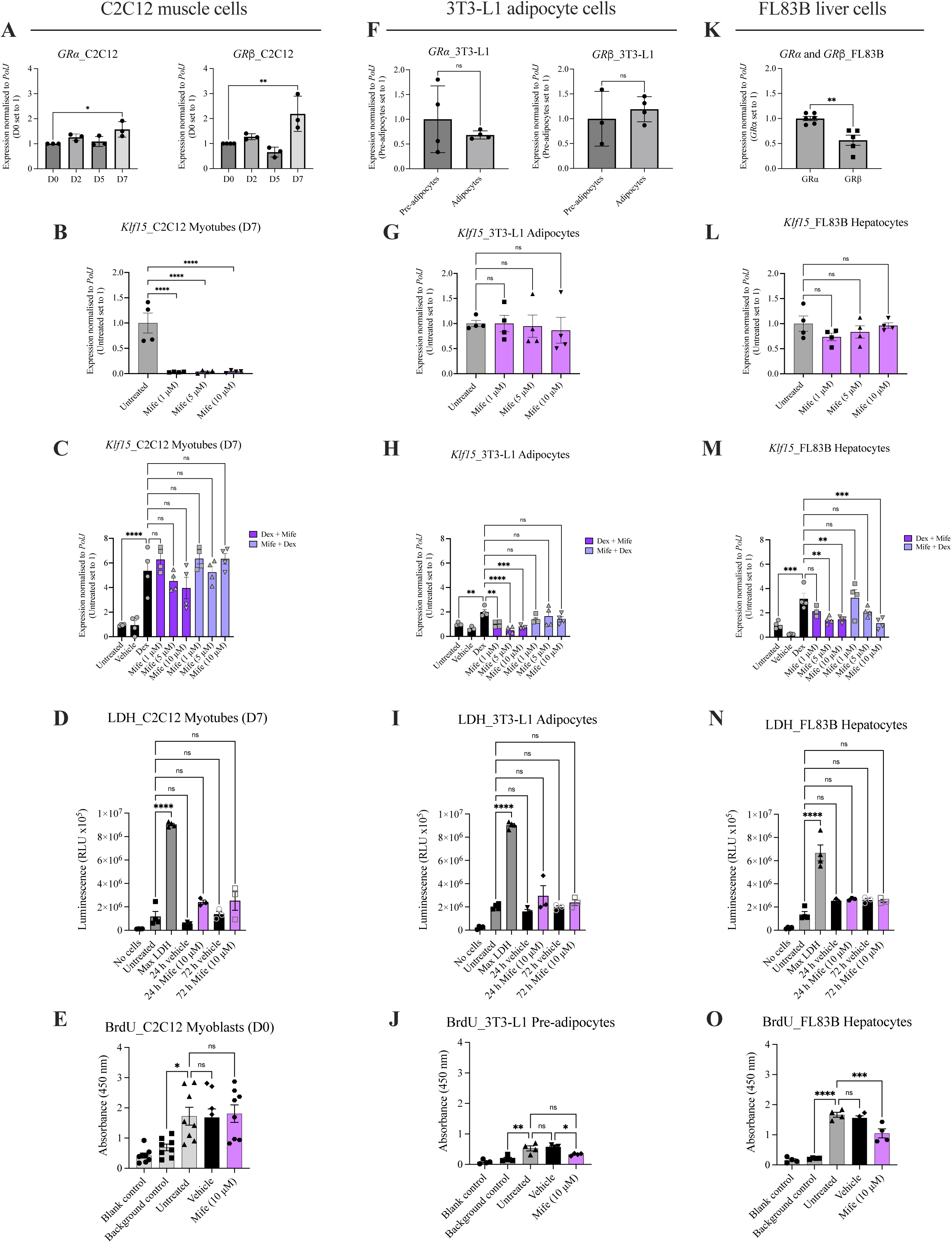
Mifepristone reduces *Klf15*-induced expression in various cellular models of metabolically active tissues. **A**, GRα and GRβ isoform expression in C2C12 cells over a 7 day (D7) differentiation period. Data are mean ± SEM, N = 3 experimental repeats (3-4 wells/repeat), one-way ANOVA, \**P*<0.05, \*\**P*<0.01. **B**, *Klf15* expression following mifepristone treatment (1, 5 or 10 μM) for 8 hours in C2C12 myotubes (D7). Untreated cells served as control. Data are mean ± SEM, N = 4 experimental repeats (3-4 wells/repeat), one-way ANOVA, \*\*\*\**P*<0.0001. **C**, *Klf15* expression following mifepristone (1, 5 or 10 μM) for 2 hours followed by dexamethasone (10 μM) for 8 hours or vice versa in C2C12 myotubes (D7). Untreated and vehicle-treated cells served as controls. Data are mean ± SEM, N = 4 experimental repeats (3-4 wells/repeat), one-way ANOVA, ns = not significant, \*\*\*\**P*<0.0001. **D**, LDH assay (cell death assay) in C2C12 myotubes (D7) following 24 hours and 72 hours treatment with mifepristone (10 μM). No cells, untreated cells and max LDH served as controls. Data are mean ± SEM, N = 3-4 experimental repeats (3-4 wells/repeat), one-way ANOVA, ns = not significant, \*\*\*\**P*<0.0001. **E**, BrdU assay (proliferation assay) in C2C12 myoblasts (D0) following 72 hours treatment with mifepristone (10 μM). Blank, background and untreated cells served as controls. Data are mean ± SEM, N = 8 experimental repeats (3-4 wells/repeat), one-way ANOVA, ns = not significant, \**P*<0.05. **F**, GRα and GRβ isoform expression in 3T3-L1 pre-adipocytes and adipocytes. Data are mean ± SEM, N = 3-4 experimental repeats (3-4 wells/repeat), unpaired *t-test*, ns = not significant. **G**, *Klf15* expression following mifepristone treatment (1, 5 or 10 μM) for 4 hours in 3T3-L1 adipocytes. Untreated cells served as control. Data are mean ± SEM, N = 4 experimental repeats (3-4 wells/repeat), one-way ANOVA, ns = not significant. **H**, *Klf15* expression following mifepristone (1, 5 or 10μM) for 2 hours followed by dexamethasone (10 μM) for 4 hours or vice versa in 3T3-L1 adipocytes. Untreated and vehicle-treated cells served as controls. Data are mean ± SEM, N = 4 experimental repeats (3-4 wells/repeat), one-way ANOVA, ns = not significant, \*\**P*<0.01, \*\*\**P*<0.001, \*\*\*\**P*<0.0001. **I**, LDH assay (cell death assay) in 3T3-L1 adipocytes following 24 hours and 72 hours treatment with mifepristone (10 μM). No cells, untreated cells and max LDH served as controls. Data are mean ± SEM, N = 3-4 experimental repeats (3-4 wells/repeat), one-way ANOVA, ns = not significant, \*\*\*\**P*<0.0001. **J**, BrdU assay (proliferation assay) in 3T3-L1 pre-adipocytes following 72 hours treatment with mifepristone (10 μM). Blank, background and untreated cells served as controls. Data are mean ± SEM, N = 4 experimental repeats (3-4 wells/repeat), one-way ANOVA, ns = not significant, \**P*<0.05, \*\**P*<0.01. **K**, GRα and GRβ isoform expression in FL83B cells. Data are mean ± SEM, N = 5-6 experimental repeats (3-4 wells/repeat), unpaired *t-test*, *P* = 0.0021. **L**, *Klf15* expression following mifepristone treatment (1, 5 or 10 μM) for 24 hours in FL83B cells. Untreated cells served as control. Data are mean ± SEM, N = 4 experimental repeats (3-4 wells/repeat), one-way ANOVA, ns = not significant. **M**, *Klf15* expression following mifepristone (1, 5 or 10 μM) for 2 hours followed by dexamethasone (10 μM) for 24 hours or vice versa in FL83B cells. Untreated and vehicle-treated cells served as controls. Data are mean ± SEM, N = 4 experimental repeats (4 wells/repeat), one-way ANOVA, ns = not significant, \*\**P*<0.01, \*\*\**P*<0.001. **N**, LDH assay (cell death assay) in FL83B cells following 24 hours and 72 hours treatment with mifepristone (10 μM). No cells, untreated cells and max LDH served as controls. Data are mean ± SEM, N = 3-4 experimental repeats (3-4 wells/repeat), one-way ANOVA, ns = not significant, \*\*\*\**P*<0.0001. **O**, BrdU assay (proliferation assay) in FL83B cells following 72 hours treatment with mifepristone (10 μM). Blank, background and untreated cells served as controls. Data are mean ± SEM, N = 4 experimental repeats (3-4 wells/repeat), one-way ANOVA, ns = not significant, \*\*\**P*<0.001, \*\*\*\**P*<0.0001.

In 3T3-L1 cells, there was no significant difference in GRα or GRβ expression between pre-adipocytes and differentiated adipocytes (Figure 1F). Mifepristone treatment alone had no effect on *Klf15* levels in differentiated 3T3-L1 adipocytes when compared to untreated cells (Figure 1G). Interestingly, all three doses of mifepristone significantly reduced *Klf15* expression when administered after dexamethasone in differentiated 3T3-L1 adipocytes when compared to dexamethasone treatment alone (Figure 1H), suggesting that mifepristone can reduce *Klf15* levels in 3T3-L1 adipocytes only when *Klf15* expression is in a hyperactivated state. We found that the highest dose of mifepristone (10 μM) did not significantly impact cell death of 3T3-L1 adipocytes when compared to untreated cells (Figure 1I). However, we did observe that the highest dose of mifepristone (10 μM) significantly reduced the proliferation of 3T3-L1 pre-adipocytes when compared to untreated cells (Figure 1J).

As FL83B hepatocyte cells are already in a differentiated state, we directly compared expression of GR isoform and found that the levels of GRβ are significantly lower than GR*α* (Figure 1K). Mifepristone treatment alone in FL83B hepatocyte cells had no effect on *Klf15* expression (Figure 1L). However, mifepristone significantly reduced *Klf15* expression at the higher doses of 5 μM and 10 μM concentrations after dexamethasone treatment when compared to dexamethasone alone (Figure 1M). The highest dose of mifepristone (10 μM) applied prior to dexamethasone also significantly reduced *Klf15* expression (Figure 1M).. Similar to 3T3-L1 cells, we found that the highest dose of mifepristone (10 μM) did not significantly impact cell death of FL83B hepatocyte cells (Figure 1N) but did significantly reduce proliferation of FL83B cells at the highest dose (Figure 1O).

Combined, our *in vitro* experiments suggest that mifepristone’s effect on *Klf15* expression, cell viability and proliferation differs between metabolic tissue cell types.

### Mifepristone treatment improves righting reflex and survival in *Smn^2B/-^*SMA mice

Having evaluated the activity of mifepristone in relevant cell types, we next wanted to assess its therapeutic potential in the *Smn^2B/-^* intermediate SMA mouse model ^39^. We first assessed the expression of *Klf15* in triceps of symptomatic (post-natal day (P) 21) *Smn^2B/-^* SMA mice and showed a significantly increased expression compared to age-matched wild type (WT) mice (Figure 2A), reproducing our previously published results ^3^. Using human primary myoblasts isolated from deltoid biopsies, we also observed significantly elevated levels of *Klf15* in myoblasts from SMA Type III patient samples compared to healthy controls (Figure 2A), again supporting our previous study reporting elevated *Klf15* expression in human SMA muscle ^3^. Both our past and current work therefore support the rationale behind reducing *Klf15* activity in SMA.

**Figure 2.**
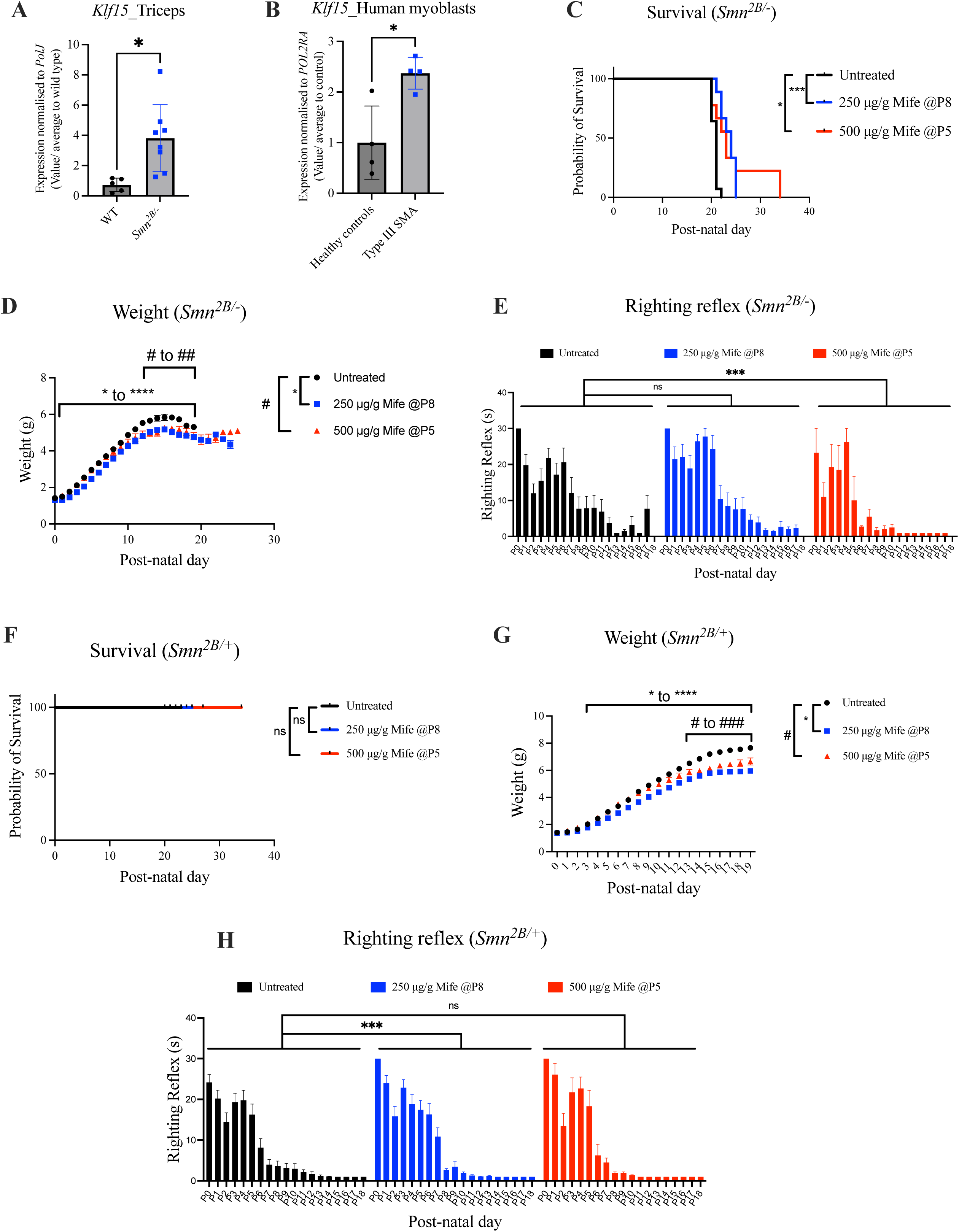
Mifepristone treatment improves disease phenotypes in *Smn^2B/-^* mice. **A**, *Klf15* expression in triceps of post-natal day (P) 21 WT and *Smn^2B/-^* mice. Data are mean ± SEM, N = 5-8 animals per experimental group, unpaired *t-test*, \**P*<0.05. **B**, *Klf15* expression in human control and SMA deltoid myoblasts, N = 4, unpaired *t-test*, \**P*<0.05. **C**, Survival curves of untreated and mifepristone-treated (250 μg/g starting at P8 or 500 μg/g starting at P5) *Smn^2B/-^* mice. Data are Kaplan-Meier survival curves, N = 9-14 animals per experimental group, Log-rank (Mantel-Cox) test, \**P*<0.05, \*\*\**P*<0.001. Survival data for selected optimal doses are repeated in supplemental figure 2A. **D**, Daily weights of untreated and mifepristone-treated (250 μg/g starting at P8 or 500 μg/g starting at P5) *Smn^2B/-^* mice. Data are mean ± SEM, N = 9-14 animals per experimental group, two-way ANOVA, *\*/#P<0.05, **/##P<0.01, ****P<0.0001.* **E**, Daily righting reflex of untreated and mifepristone-treated (250 μg/g starting at P8 or 500 μg/g starting at P5) *Smn^2B/-^* mice. Data are mean ± SEM, N = 9-14 animals per experimental group, one-way ANOVA, ns = not significant, \*\*\**P*<0.001. **F**, Survival curves of untreated and mifepristone-treated (250 μg/g starting at P8 or 500 μg/g starting at P5) *Smn^2B/+^* mice. Data are Kaplan-Meier survival curves, N = 11-14 animals per experimental group, Log-rank (Mantel-Cox) test, ns = not significant. **G**, Daily weights of untreated and mifepristone-treated (250 μg/g starting at P8 or 500 μg/g starting at P5) *Smn^2B/+^*mice. Data are mean ± SEM, N = 11-14 animals per experimental group, two-way ANOVA, *\*/#P<0.05, **/##P<0.01, ***/###P<0.001, ****P<0.0001.* H, Daily righting reflex of untreated and mifepristone-treated (250 μg/g starting at P8 or 500 μg/g starting at P5) *Smn^2B/-^* mice. Data are mean ± SEM, N = 11-14 animals per experimental group, one-way ANOVA, ns = not significant, \*\*\**P*<0.001.

We initially conducted pilot studies to optimise the dosing regimen of mifepristone in *Smn^2B/-^*mice, which determined that the best dosing regimens were 500 μg/g daily gavage starting at P5 and 250 μg/g daily gavage starting at P8 (Supplementary Figure 2A). We also found that the vehicle (0.5% carboxymethylcelluose (CMC)) did not affect weight, righting reflex or survival of *Smn^2B/-^* SMA mice and *Smn^2B^*^/+^ healthy littermates when compared to untreated animals (Supplementary Figure 2B-G).

As demonstrated in the pilot studies, both optimal mifepristone dosing regimens (500 μg/g daily gavage starting at P5 and 250 μg/g daily gavage starting at P8) significantly increased the lifespan of *Smn^2B/-^* SMA mice compared to untreated *Smn^2B/-^*animals (Figure 2C), of which 500 μg/g improved life expectancy for longer. In contrast, the weights of *Smn^2B/-^* SMA mice were significantly decreased across both mifepristone-treated compared to untreated *Smn^2B/-^* mice (Figure 2D). Of note, this reduced weight occurred prior to treatment (P8) in the 250 μg/g experimental group suggesting that this may be due to smaller weights at birth in those treated litters. We also saw a significant decrease in the time it took 500 μg/g mifepristone-treated *Smn^2B/-^* mice to right themselves during disease progression compared to untreated *Smn^2B/-^* mice (Figure 2E). However, there was no significant difference in the righting reflex between 250 μg/g mifepristone-treated and untreated animals (Figure 2E), suggesting that timing and dose of mifepristone lead to differential effects in *Smn^2B/-^* mice. The lifespan of *Smn^2B/+^*healthy control mice was not negatively affected by mifepristone (Figure 2F). However, similar to *Smn^2B/-^* mice, both dosing regimens of mifepristone significantly decreased the weight of *Smn^2B/+^* animals (Figure 2G), which preceded the first dose (P8) in 250 μg/g mifepristone-treated litters, further supporting intrinsic smaller weights at birth in that experimental cohort. Interestingly, the time to right in *Smn^2B/+^* healthy littermates was significantly decreased following 250 μg/g mifepristone treatment, while there was no impact on righting reflex in 500 μg/g mifepristone-treated *Smn^2B/+^* mice when compared to untreated *Smn^2B/+^*animals (Figure 2H). As 250 μg/g mifepristone was administered at a later time point this could suggest that mifepristone’s effect in healthy control mice is treatment stage dependent.

### Mifepristone treatment is well tolerated but ineffective in the severe ‘Taiwanese’ SMA mouse model

As there is a wide range of SMA mouse models with differing disease onsets and progression, we next wanted to investigate mifepristone treatment in a more severe mouse model. We treated severe Taiwanese *Smn^-/-^;SMN2* mice ^40^ with mifepristone daily starting at P3. An earlier start date (P3 vs P5 in *Smn^2B/-^* mice) was selected due to the earlier and more severe disease onset in these mice. We found that there was no significant differences in survival, weight and righting reflex between vehicle-treated and mifepristone-treated *Smn^-/-^;SMN2* mice compared to untreated animals (Supplementary Figure 3A-C). In addition, no adverse effects were observed in mifepristone-treated *Smn^+/-^;SMN2* healthy littermates compared to untreated and vehicle-treated animals (Supplementary Figure 3D-F). Overall, our results highlight key differences between SMA mouse models in their response to mifepristone, specifically demonstrating the more aggressive nature of phenotypes in the Taiwanese model making it much harder to rescue. This is in line with previous studies demonstrating differential therapeutic effects of interventions in different SMA mouse models ^3,41^.

### Mifepristone treatment ameliorates neuromuscular pathology in a severe *C. elegans* model of SMA

Next, we investigated mifepristone in a validated severe SMA *C. elegans* invertebrate model ^42,43^. The *C. elegans* nematode maintains functional conservation of neuronal processes and has high homology with the human genome ^44^. *C. elegans* have a single *smn-1* gene that, when diminished causes larval lethality, slowed growth and impaired neuromuscular function in pharyngeal pumping and mobility ^33^. Here, SMA *C.elegans smn-1* (ok355) and control *C.elegans smn-1/hT2* nematodes were treated with increasing doses of mifepristone (1, 15 and 30 μM) and compared to vehicle. Interestingly, we observed that higher 15 and 30 μM concentrations of mifepristone significantly increased mobility forward time in *C.elegans smn-1* (ok355) compared to vehicle-treated *C.elegans smn-1* (ok355) (Figure 3A). Additionally, the highest dose of mifepristone (30 μM) significantly increased pharyngeal pumping in the *C.elegans smn-1* (ok355) model compared to vehicle treated *C.elegans smn-1* (ok355) (Figure 3B). Interestingly, mifepristone did not impact mobility forward time or pharyngeal pumping in the *C.elegans smn-1/hT2* controls (Figure 3C-D), suggesting a disease specific effect of mifepristone on neuromuscular function in nematodes. Of note, other parameters such as distance travelled, speed and reversal times remained unchanged in both SMA *C.elegans smn-1* (ok355) and control *C.elegans smn-1/hT2* nematodes (Supplementary Figure 4).

**Figure 3.**
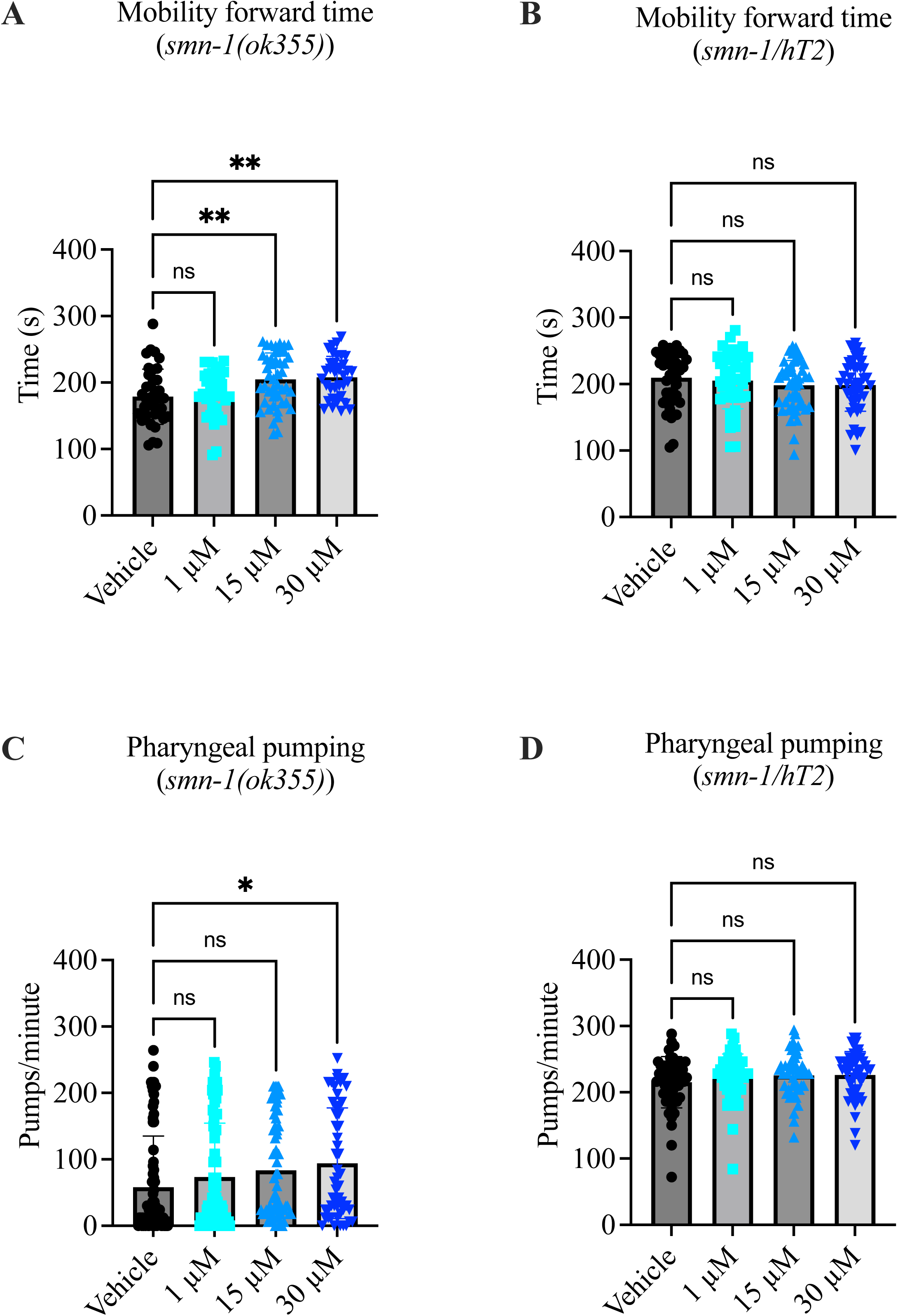
Mifepristone ameliorates the neuromuscular phenotype in a severe SMA *C. elegans smn-1* (ok355) model. **A**, Mobility forward time filmed at 15 frames/second for 5 minutes in vehicle or mifepristone-treated (1, 15 or 30 μM) SMA *C. elegans smn-1* (ok355). Data are mean ± SEM, N = 25 animals per experimental group, one-way ANOVA, ns = not significant, *\*\*P*<0.01. **B**, Mobility forward time filmed at 15 frames/ second for 5 minutes in vehicle or mifepristone-treated (1, 15 or 30 μM) control *C. elegans smn-1/hT2*. Data are mean ± SEM, N = 25 animals per experimental group, one-way ANOVA, ns = not significant. **C**, Pharyngeal pumping rates (pumps/minute) defined as grinder movements in any axis at 175 frames/10 seconds in vehicle or mifepristone-treated (1, 15 or 30 μM) SMA *C. elegans smn-1* (ok355). Data are mean ± SEM, N = 25 animals per experimental group, one-way ANOVA, ns = not significant, \**P*<0.05. **D**, Pharyngeal pumping rates (pumps/minute) defined as grinder movements in any axis at 175 frames/10 seconds in vehicle or mifepristone-treated (1, 15 or 30 μM) control *C. elegans smn-1/hT2*. One-way ANOVA was performed. Data are mean ± SEM, N = 25 animals per experimental group, one-way ANOVA, ns = not significant.

Our data therefore support the beneficial effects of mifepristone in two different models of SMA.

### Mifepristone significantly downregulates the expression of *Klf15* in BAT and increases myofiber area in skeletal muscle of *Smn^2B/-^* mice

Next, we wanted to investigate the effect of mifepristone at molecular and histological levels in metabolically active skeletal muscle, liver and BAT of *Smn^2B/-^* SMA and *Smn^2B/+^*control mice. Firstly, we investigated *Klf15* expression in tissues from P18 untreated, 250 μg/g (P8) and 500 μg/g (P5) mifepristone-treated mice. In *Smn^2B/-^* mice, we observed that *Klf15* expression in liver and triceps remained unchanged between untreated and mifepristone-treated *Smn^2B/-^* animals (Figure 4A-B). However, both doses of mifepristone significantly reduced *Klf15* expression in BAT from *Smn^2B/-^* mice compared to untreated *Smn^2B/-^* animals (Figure 4C). Similar results were found in liver, triceps and BAT of *Smn^2B/+^* mice (Figure 4D-F), suggesting increased activity of the GR antagonist in adipose tissue, likely due to the increased metabolic rate of BAT.

**Figure 4.**
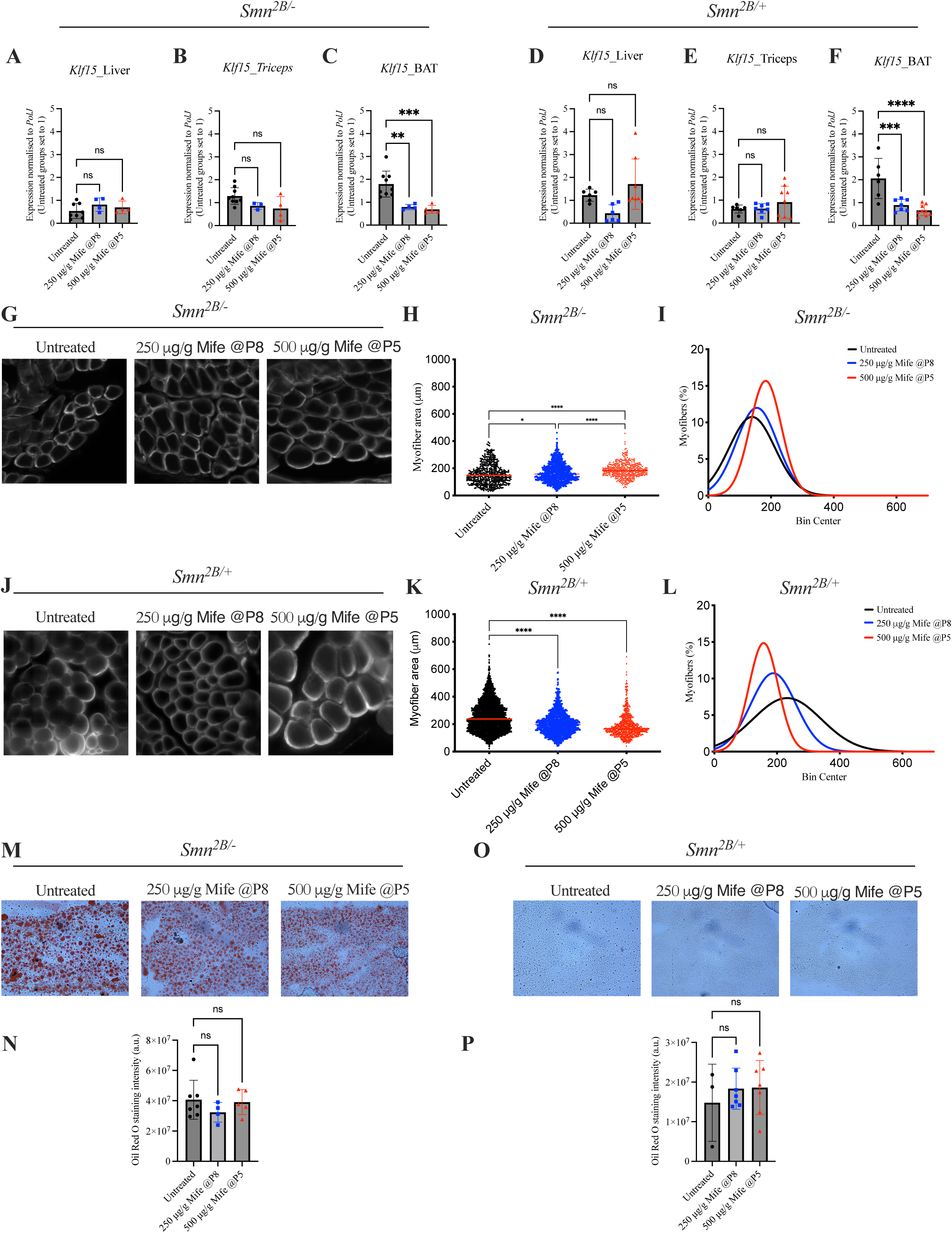
Mifepristone impacts disease phenotype in *Smn^2B/-^* mice in a tissue and disease-state specific manner. **A**, *Klf15* expression in liver from post-natal day (P) 18 untreated and mifepristone-treated (250 μg/g starting at P8 or 500 μg/g starting at P5) *Smn^2B/-^* mice. Data are mean ± SEM, N = 4-8 animals per experimental group, one-way ANOVA, ns = not significant. **B**, *Klf15* expression in triceps from P18 untreated and mifepristone-treated (250 μg/g starting at P8 or 500 μg/g starting at P5) *Smn^2B/-^* mice. Data are mean ± SEM, N = 3-8 animals per experimental group, one-way ANOVA, ns = not significant. **C**, *Klf15* expression in brown adipose tissue (BAT) from P18 untreated and mifepristone-treated (250 μg/g starting at P8 or 500 μg/g starting at P5) *Smn^2B/-^* mice. Data are mean ± SEM, N = 4-8 animals per experimental group, one-way ANOVA, \*\**P*<0.01, \*\*\**P*<0.001. **D**, *Klf15* expression in liver from P18 untreated and mifepristone-treated (250 μg/g starting at P8 or 500 μg/g starting at P5) *Smn^2B/+^* mice. Data are mean ± SEM, N = 6-8 animals per experimental group, one-way ANOVA, ns = not significant. **E**, *Klf15* expression in triceps from P18 untreated and mifepristone-treated (250 μg/g starting at P8 or 500 μg/g starting at P5) *Smn^2B/+^* mice. Data are mean ± SEM, N = 6-9 animals per experimental group, one-way ANOVA, ns = not significant. **F**, *Klf15* expression in BAT from P18 untreated and mifepristone-treated (250 μg/g starting at P8 or 500 μg/g starting at P5) *Smn^2B/+^* mice. Data are mean ± SEM, N = 6-9 animals per experimental group, one-way ANOVA, \*\*\**P*<0.001, \*\*\*\**P*<0.0001. G, Representative images of laminin-stained cross-sections of tibialis anterior (TA) muscles from P18 untreated and mifepristone-treated (250 μg/g starting at P8 or 500 μg/g starting at P5) *Smn^2B/-^* mice. **H**, Quantification of myofiber area of laminin-stained cross-sections of TA muscles from P18 untreated and mifepristone-treated (250 μg/g starting at P8 or 500 μg/g starting at P5) *Smn^2B/-^* mice. Data are dot plot and mean, n = 3-4 animals per experimental group (>200 myofibers per experimental group), one-way ANOVA, \**P*<0.05, \*\*\*\**P*<0.0001. **I**, Relative frequency distribution of myofiber size in TA muscles from P18 untreated and mifepristone-treated (250 μg/g starting at P8 or 500 μg/g starting at P5) *Smn^2B/-^* mice. **J**, Representative images of laminin-stained cross-sections of TA muscles from P18 untreated and mifepristone-treated (250 μg/g starting at P8 or 500 μg/g starting at P5) *Smn^2B/+^*mice. **K**, Quantification of myofiber area of laminin-stained cross-sections of TA muscles from P18 untreated and mifepristone-treated (250 μg/g starting at P8 or 500 μg/g starting at P5) *Smn^2B/+^* mice. Data are dot plot and mean, n = 3-4 animals per experimental group (>200 myofibers per experimental group), one-way ANOVA, \*\*\*\**P*<0.0001. **L**, Relative frequency distribution of myofiber size in TA muscles from P18 untreated and mifepristone-treated (250 μg/g starting at P8 or 500 μg/g starting at P5) *Smn^2B/+^* mice. **M**, Representative images of oil red O-stained liver sections from P18 untreated and mifepristone-treated (250 μg/g starting at P8 or 500 μg/g starting at P5) *Smn^2B/-^* mice. **N**, Quantification of oil red O staining intensity in liver sections P18 untreated and mifepristone-treated (250 μg/g starting at P8 or 500 μg/g starting at P5) *Smn^2B/-^* mice. Data are mean ± SEM, N = 4-7 animals per experimental group, one-way ANOVA, ns = not significant. **O**, Representative images of oil red O-stained liver sections from P18 untreated and mifepristone-treated (250 μg/g starting at P8 or 500 μg/g starting at P5) *Smn^2B/+^* mice. **P**, Quantification of oil red O staining intensity in liver sections P18 untreated and mifepristone-treated (250 μg/g starting at P8 or 500 μg/g starting at P5) *Smn^2B/+^* mice. Data are mean ± SEM, N = 3-7 animals per experimental group, one-way ANOVA, ns = not significant.

As muscle atrophy is a canonical pathology of SMA, we next analysed myofiber area of the *tibialis anterior* (TA) muscle from P18 untreated, 250 μg/g and 500 μg/g mifepristone-treated *Smn^2B/-^* and *Smn^2B/+^* mice. Treatment with both doses of mifepristone significantly increased myofiber size in mifepristone-treated *Smn^2B/-^*mice when compared to untreated *Smn^2B/-^* animals (Figure 4G-I). Interestingly, myofiber area in *Smn^2B/+^* healthy littermate controls was significantly decreased following 250 μg/g and 500 μg/g mifepristone treatment when compared to untreated *Smn^2B/+^* animals (Figure 4J-L), which may explain the reduction in body weight that is seen following mifepristone treatment in *Smn^2B/+^* mice.

Finally, mifepristone has been successfully used to treat fatty liver disease in patients with CS ^45^, therefore, we used an oil-red-O stain to assess whether mifepristone affected the previously reported lipid accumulation in the liver of *Smn^2B/-^* mice ^10^. Quantification of oil-red-O staining intensity showed no significant differences between untreated *Smn^2B/-^* mice and mifepristone-treatment *Smn^2B/-^*animals (Figure 4M-N). Similar results were observed in *Smn^2B/+^* mice where there is a noticeable absence of hepatic lipid accumulation (Figure 4O-P).

Overall, our molecular and histological analyses show tissue- and disease state-specific effects of mifepristone on skeletal muscle, liver and BAT.

### Combinatorial treatment of scAAV9-*SMN1* with 500 μg/g mifepristone leads to selective synergistic effects in *Smn^2B/-^* SMA mice

To address the benefit of combinatorial therapy for SMA, we combined daily administration of mifepristone (from P5 to P21) with an Onasemnogene abeparvovec-like vector (scAAV9-*SMN1*) ^46^ given at P0 (day of birth) via facial intravenous injection (IV) with a scAAV9-*GFP* vector used as a control. Even though both doses of mifepristone improved survival of *Smn^2B/-^* mice, 22% of mice survived an additional 10 days longer (to P35) following 500 μg/g compared to 250 μg/g mifepristone. Mifepristone (500 μg/g) produced a substantially greater improvement in motor function and myofiber hypertrophy than the alternative dose, without any adverse effects in healthy littermate controls. Furthermore, 500 μg/g mifepristone was administered at the earlier age of P5 (in contrast to 250 μg/g mifepristone (P8)), a translational time point likely to produce greater therapeutic outcome and correspond with administration to human patients with SMA. Taken together we moved forward with 500 μg/g mifepristone to assess combinatorial therapy.

We assessed phenotypic outcomes, including weight, righting reflex and survival in *Smn^2B/-^* and *Smn^2B/+^* mice. Daily weights were recorded from birth (P0) until P28 and weekly weights were recorded from P28 onwards. Of note, as scAAV9-*GFP*-injected *Smn^2B/-^* SMA mice did not survive past weaning (P21), it was only possible to collect data from scAAV9-*SMN1* and scAAV9-*SMN1* + 500 μg/g mifepristone animals. In pre-weaned *Smn^2B/-^* animals, we found no significant differences between scAAV9-*SMN1*-injected animals compared to scAAV9-*SMN1* + 500 μg/g mifepristone (Supplementary Figure 5A). However, scAAV9-*GFP*-injected *Smn^2B/-^* SMA mice weighed significantly less than scAAV9-*SMN1*-injected *Smn^2B/-^* SMA mice and the scAAV9-*SMN1* + 500 μg/g mifepristone *Smn^2B/-^*SMA mice as disease progressed, due to the lack of therapeutic activity of this control vector (Supplementary Figure 5A). Interestingly, while the righting reflex of scAAV9-*SMN1*-injected *Smn^2B/-^* animals was similar to those treated with scAAV9-*SMN1* + 500 μg/g mifepristone, there was a significant improvement of the righting reflex in scAAV9-*SMN1*-injected *Smn^2B/-^* SMA mice compared to the control vector, scAAV9-*GFP*-injected *Smn^2B/-^* SMA mice (Supplementary Figure 5B). As weight differences become noticeable between the sexes post-weaning, we separated male and female mice from beyond this time-point. We observed no significant difference in weight between scAAV9-*SMN1*-injected *Smn^2B/-^* males and females compared to males and females treated with scAAV9-*SMN1* + 500 μg/g mifepristone from P28 to humane endpoint (Supplementary Figure 5C-D). Similar analyses in *Smn^2B/+^* healthy control littermates revealed comparable weights and righting reflex between scAAV9-*GFP*-injected, scAAV9-*SMN1*-injected and those treated with scAAV9-*SMN1* + 500 μg/g mifepristone (Supplementary Figure 5E-H).

We also undertook molecular and histological investigations in tissues from 6-month-old *Smn^2B/-^* SMA mice and *Smn^2B/+^* control mice that either received a single injection of scAAV9-*SMN1* or were treated with the combinatorial intervention of scAAV9-*SMN1* + 500 μg/g mifepristone. We assessed *Klf15* expression in liver, skeletal muscle (triceps) and BAT. In *Smn^2B/-^* SMA mice *Klf15* expression was significantly reduced in the liver (Figure 5A), while remaining unchanged in the triceps (Figure 5B) and BAT (Figure 5C) of animals treated with scAAV9-*SMN1* + 500 μg/g mifepristone compared to scAAV9-*SMN1* alone. In *Smn^2B/+^* control mice, *Klf15* expression in liver (Figure 5D) and triceps (Figure 5E) remained unchanged in animals treated with scAAV9-*SMN1* + 500 μg/g mifepristone compared to scAAV9-*SMN1* alone, whereas there was a significant increase of *Klf15* expression in BAT following scAAV9-*SMN1* + 500 μg/g mifepristone when compared to scAAV9-*SMN1* alone (Figure 5F). Our results therefore suggest disease state and tissue-specific effect the combinatorial intervention on *Klf15* expression.

**Figure 5.**
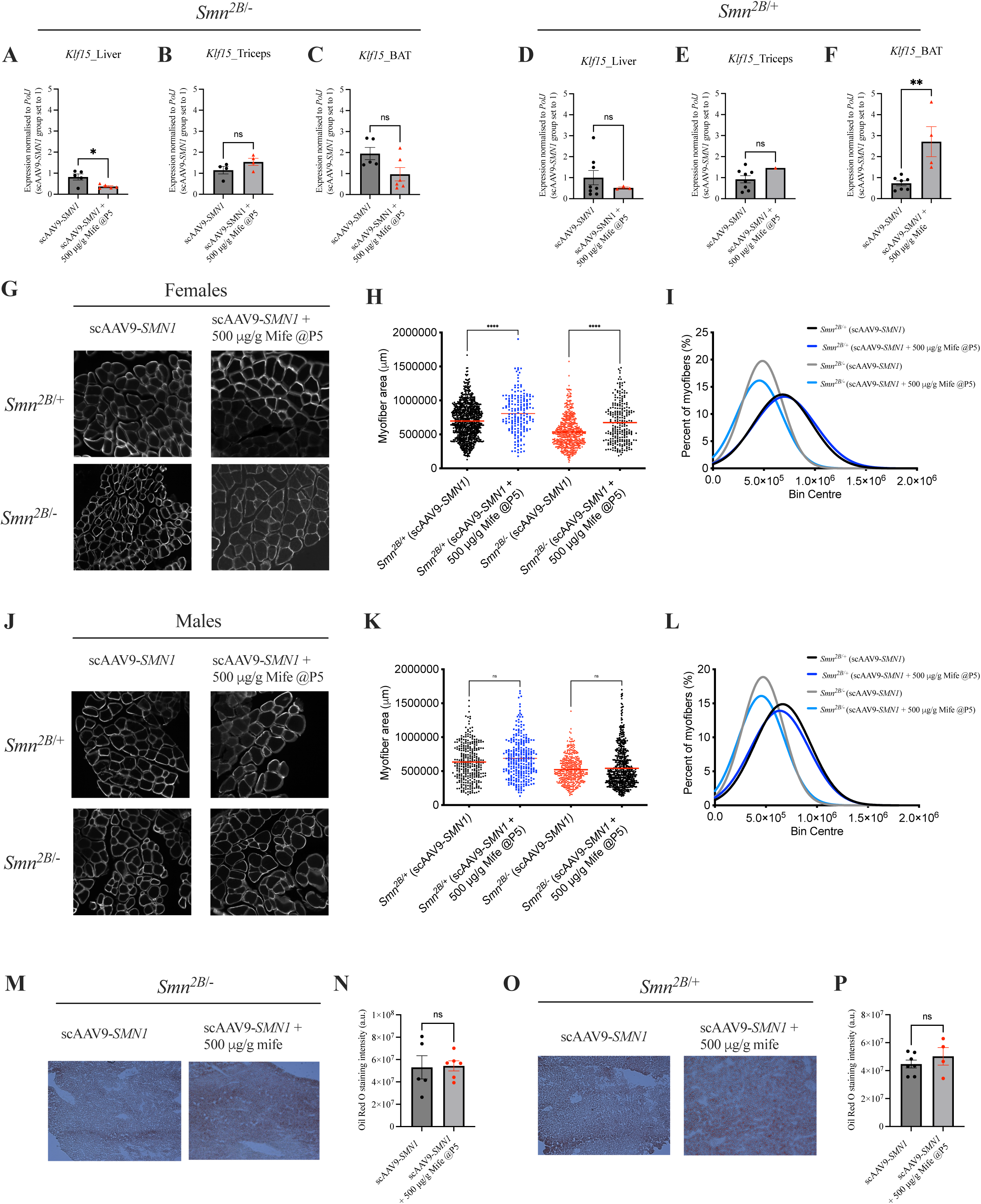
Combinatorial treatment of scAAV9-*SMN1* and mifepristone reduces *Klf15* expression in the liver of *Smn^2B/-^* mice and significantly increases myofiber area in females specifically. **A**, *Klf15* expression in liver from 6-month-old *Smn^2B/-^* mice following treatment with scAAV9-*SMN1* at post-natal day (P) 0 or a combination of scAAV9-*SMN1* + mifepristone (500 μg/g from P5-P21). Data are mean ± SEM, N = 5 animals per experimental group, unpaired *t-test*, \**P*<0.05. **B**, *Klf15* expression in triceps from 6-month-old *Smn^2B/-^* mice following treatment with scAAV9-*SMN1* at post-natal P0 or a combination of scAAV9-*SMN1* + mifepristone (500 μg/g from P5-P21). Data are mean ± SEM, N = 4 animals per experimental group, unpaired *t-test*, ns = not significant. **C**, *Klf15* expression in brown adipose tissue (BAT) from 6-month-old *Smn^2B/-^* mice following treatment with scAAV9-*SMN1* at post-natal day (P) 0 or a combination of scAAV9-*SMN1* + mifepristone (500 μg/g from P5-P21). Data are mean ± SEM, N = 5 animals per experimental group, unpaired *t-test*, ns = not significant. **D**, *Klf15* expression in liver from 6-month-old *Smn^2B/+^* mice following treatment with scAAV9-*SMN1* at post-natal P0 or a combination of scAAV9-*SMN1* + mifepristone (500 μg/g from P5-P21). Data are mean ± SEM, N = 4-8 animals per experimental group, unpaired *t-test*, ns = not significant. **E**, *Klf15* expression in triceps from 6-month-old *Smn^2B/+^* mice following treatment with scAAV9-*SMN1* at post-natal P0 or a combination of scAAV9-*SMN1* and mifepristone (500 μg/g from P5-P21). Data are mean ± SEM, N = 4-8 animals per experimental group, unpaired *t-test*, ns = not significant. **F**, *Klf15* expression in BAT from 6-month-old *Smn^2B/+^* mice following treatment with scAAV9-*SMN1* at post-natal P0 or a combination of scAAV9-*SMN1* + mifepristone (500 μg/g from P5-P21). Data are mean ± SEM, N = 4-8 animals per experimental group, unpaired *t-test*, \*\**P*<0.01. **G**, Representative images of laminin-stained cross-sections of *tibialis anterior* (TA) muscles from 6-month-old *Smn^2B/-^* and *Smn^2B/+^* females following treatment with scAAV9-*SMN1* at post-natal P0 or a combination of scAAV9-*SMN1* + mifepristone (500 μg/g from P5-P21). **H**, Quantification of myofiber area of laminin-stained cross-sections of TA muscles from 6-month-old *Smn^2B/-^* and *Smn^2B/+^* females following treatment with scAAV9-*SMN1* at post-natal P0 or a combination of scAAV9-*SMN1* + mifepristone (500 μg/g from P5-P21). Data are dot plot and mean, n = 3-5 animals per experimental group (>200 myofibers per experimental group), one-way ANOVA, \*\*\*\**P*<0.0001. **I**, Relative frequency distribution of myofiber size in TA muscles from 6-month-old *Smn^2B/-^* and *Smn^2B/+^* females following treatment with scAAV9-*SMN1* at post-natal P0 or a combination of scAAV9-*SMN1* + mifepristone (500 μg/g from P5-P21). **J**, Representative images of laminin-stained cross-sections of TA muscles from 6-month-old *Smn^2B/-^* and *Smn^2B/+^* males following treatment with scAAV9-*SMN1* at post-natal P0 or a combination of scAAV9-*SMN1* + mifepristone (500 μg/g from P5-P21). **K**, Quantification of myofiber area of laminin-stained cross-sections of TA muscles from 6-month-old *Smn^2B/-^* and *Smn^2B/+^* males following treatment with scAAV9-*SMN1* at post-natal P0 or a combination of scAAV9-*SMN1* and mifepristone (500 μg/g from P5-P21). Data are dot plot and mean, n = 3-5 animals per experimental group (>200 myofibers per experimental group), one-way ANOVA, ns = not significant. **L**, Relative frequency distribution of myofiber size in TA muscles from 6-month-old *Smn^2B/-^* and *Smn^2B/+^* males following treatment with scAAV9-*SMN1* at post-natal P0 or a combination of scAAV9-*SMN1* + mifepristone (500 μg/g from P5-P21). **M**, Representative images of oil red O-stained liver sections from 6-month-old *Smn^2B/-^* mice following treatment with scAAV9-*SMN1* at post-natal P0 or a combination of scAAV9-*SMN1* + mifepristone (500 μg/g from P5-P21). **N**, Quantification of oil red O staining intensity in liver sections from 6-month-old *Smn^2B/-^* mice following treatment with scAAV9-*SMN1* at post-natal P0 or a combination of scAAV9-*SMN1* + mifepristone (500 μg/g from P5-P21). Data are mean ± SEM, N = 5-6 animals per experimental group, unpaired *t-test*, ns = not significant. **O**, Representative images of oil red O-stained liver sections from 6-month-old *Smn^2B/+^* mice following treatment with scAAV9-*SMN1* at post-natal P0 or a combination of scAAV9-*SMN1* + mifepristone (500 μg/g from P5-P21). **P**, Quantification of oil red O staining intensity in liver sections from 6-month-old *Smn^2B/+^* mice following treatment with scAAV9-*SMN1* at post-natal P0 or a combination of scAAV9-*SMN1* + mifepristone (500 μg/g from P5-P21). Data are mean ± SEM, N = 4-6 animals per experimental group, unpaired *t-test*, ns = not significant.

Next, we assessed myofiber area, whereby the analyses were separated by sex. We observed that the combination treatment of scAAV9-*SMN1*+ 500 μg/g mifepristone significantly increased the myofiber area of both *Smn^2B/-^*SMA mice and *Smn^2B/+^* control females compared to scAAV9-*SMN1* alone (Figure 5G-I). In contrast, there were not significant differences in the myofiber area between treatment groups in males (Figure 5J-L), suggesting a sex-dependent effect of the combinatorial approach with mifepristone as an adjunct therapy.

Finally, we investigated whether scAAV9-*SMN1* +500 μg/g mifepristone impacted lipid accumulation in the liver, as assessed via oil red O staining intensity. Lipid accumulation remained unchanged in liver of both *Smn^2B/-^* SMA mice and *Smn^2B/+^* control mice treated with either scAAV9-*SMN1* or scAAV9-*SMN1* +500 μg/g mifepristone (Figure 5M-P), supporting previous research demonstrating rescue of the fatty liver phenotype by scAAV9-*SMN1* therapy ^46^.

## DISCUSSION

In this study, we addressed two key research questions. Firstly, whether targeting the GC-GR-*Klf15* metabolic pathway via mifepristone could improve disease phenotypes in SMA animal models ^3^. Secondly, whether combining an *SMN1*-directed gene therapy with mifepristone could lead to additional benefits in peripheral and metabolic tissues compared to the gene therapy alone. Overall, we found that repurposing mifepristone for the treatment of SMA reduced *Klf15* expression specifically in BAT, increased the lifespan and muscle size of treated *Smn^2B/-^* mice, and improved motor function in a SMA *C. elegans* model. Furthermore, combining mifepristone with an scAAV9-*SMN1* gene therapy resulted in tissue-, sex and disease state-specific improved pathological changes compared to the gene therapy alone.

In the first part of this study, we evaluated mifepristone activity in cell models of peripheral tissues. Across all three models (C2C12 (muscle), 3T3-L1 (adipose) and FL83B (liver)), treatment with mifepristone reduced dexamethasone-induced *Klf15* expression in a cell and differentiation state-dependent manner. Key players in mifepristone’s mode of action, are the isoforms of the GR (α and β), whereby GRα is the main mediator of GCs while GRβ inhibits GRα and induces GC resistance ^37,38^. We found that expression of both GR isoforms (α and β) were increased in untreated C2C12 myotubes compared to C2C12 myoblasts, suggesting that the concomitant upregulation of GRβ in myotubes may counteract the aberrant activation of GRα. Previous research overexpressing GRβ (GRβOE) in C2C12 cells showed that GC-induced leucine zipper (GILZ), a target of GRα, was significantly reduced in GRβOE myoblasts, demonstrating a reduction in GRα activity when GRβ expression is elevated ^41^. This could explain the inability of mifepristone to antagonise GRα and reduce dexamethasone-induced *Klf15* expression in our treated C2C12 myotubes. Mifepristone’s inhibition of *Klf15* expression may therefore work better when given early during muscle development.

Interestingly, in FL83B hepatocyte-like cells, mifepristone reduced *Klf15* expression regardless of dexamethasone treatment, which might be explained by the observation that FL83B cells displayed higher levels of GRα than GRβ in a differentiated state. Lower levels of GRβ suggests that mifepristone is able to actively bind GRα and antagonise transcriptional regulation without resistance from GRβ’s negative regulation ^47^. In fact, previous research has shown that small GR antagonist molecules that are active in the liver can successfully ameliorate metabolic syndrome in rats ^48^, suggesting that modulation of the GR pathway could help reduce known metabolic liver pathology in SMA ^12,19^.

A key finding from our molecular analyses of mifepristone-treated tissues is the selective downregulation of *Klf15* expression in BAT of both *Smn^2B/-^* SMA mice and *Smn^2B/+^* healthy littermate controls. The previously reported increased expression of *Klf15* in BAT of SMA mice ^3^ may disrupt *Klf15*’s ability to maintain crucial processes such as lipid metabolism. Specifically, *Klf15* has been demonstrated to regulate fuel switching between glucose and fatty acids in response to changes in energy status in BAT ^49^. The imbalance of *Klf15* levels in BAT of SMA mice may therefore contribute to aberrant gluconeogenesis and insulin resistance as a result of reduced metabolic flexibility ^49^. Reducing *Klf15* expression in BAT with mifepristone may potentially improve metabolic homeostasis in that tissue. Additionally, cross-talk occurs between BAT and pancreatic cells in obese mice, whereby the overexpression of *Klf15* in BAT from obese mice enhanced insulin secretion from pancreatic

β-cells ^50^. As previous studies have reported a particular reduction in the pancreatic β-cells in *Smn^2B/-^* mice ^13^, we speculate that the increased expression of *Klf15* seen in BAT of SMA mice could be a compensatory mechanism due to the reduced activity of pancreatic β-cells.^13^ Mifepristone’s ability to reduce *Klf15* in BAT of *Smn^2B/-^* mice could thus improve insulin resistance and metabolic pathologies through crosstalk between adipose and pancreatic tissues ^51^.

Our *in vivo* studies also revealed a significant increase in myofiber area of *Smn^2B/-^* SMA mice following mifepristone treatment. This increase in myofiber size is likely associated with the improved righting reflex times seen after mifepristone treatment (specifically 500 μg/g at P5) in *Smn^2B/-^* mice. *Klf15* plays a role in muscle physiology and exercise adaptation by regulating lipid flux, which could explain our current findings ^25,52^. Furthermore, the knockdown of DDIT4 (REDD1), another GC regulated protein, boosts muscle mass supporting the concept that targeting GC pathways as an adjunct therapy may be beneficial for SMA ^53,54^.

Our *in vivo* work was conducted in both vertebrate and invertebrate models of SMA, which led to several key observations such as the ability of mifepristone to improve the survival of *Smn^2B/-^* mice as well as improve neuromuscular pathology in both *Smn^2B/-^* mice and *C.elegans smn-1*(ok355) models. Our findings support previous work, from our group and others, that demonstrate that SMN-independent therapeutics alone can improve disease phenotypes, including lifespan ^3,20,55^. In the current study, these improvements were seen when mifepristone was administered at the later time-points of P5 and P8 in *Smn^2B/-^* mice, demonstrating that SMN-independent treatments can be administered later on during disease progression and still be able to attenuate pathology ^56^. However, mifepristone did not improve the survival of severe Taiwanese mice, suggesting that peripheral pathologies may have a greater impact on disease phenotype in milder forms of SMA ^56^. The adult population or patients with milder forms of SMA, including those now receiving therapy, may therefore benefit greatly from second-generation therapies ^57^.

Given that a single dose of scAAV9-*SMN1* can significantly increase life expectancy, we combined scAAV9-*SMN1* with 500 μg/g mifepristone in *Smn^2B/-^* mice ^58^. While delivery of *SMN1* is sufficient to significantly improve lifespan in both mouse models and patients, this therapeutic approach does have limitations ^46,59^. Indeed, as the longevity of this therapy is not yet known, it is possible that *SMN1* cDNA will become diluted over time in dividing cells, thus preventing prolonged peripheral expression ^60^. Furthermore, the amount of viral vector that be delivered is currently limited due to the viral affinity for the liver ^60^. In fact, research has shown mifepristone can control expression of transgenes that are potentially toxic, as an activator in drug-dependent inducible systems ^61^. Ultimately, there is wide clinical evidence demonstrating the safety of mifepristone following both acute and chronic dosing regimens and most patients will likely benefit from a combinatorial approach to their treatment ^60,62,63^.

In the current study, a combinatorial approach enabled us to determine any synergistic activities as well as evaluate whether we could expand the therapeutic benefit of scAAV9-*SMN1* to further improve pathology by targeting metabolic perturbations in *Smn^2B/-^* mice. While scAAV9-*SMN1* combined with 500 μg/g mifepristone did not change the phenotypic outcomes of *Smn^2B/-^* mice compared to gene therapy alone, molecular analyses of *Klf15* expression in tissues revealed differences between mifepristone-treated *Smn^2B/-^* and *Smn^2B/+^* mice. For instance, *Klf15* levels were significantly reduced in the liver of scAAV9-*SMN1* combined with 500 μg/g mifepristone-treated *Smn^2B/-^*mice, but not *Smn^2B/+^* animals that received the same combinatorial treatment. This suggests that mifepristone may preferentially target the liver when combined with an *SMN* rescuing therapy in a diseased state. As *Klf15* has been reported as a regulator of hepatic maturation, the decrease of *Klf15* in *Smn^2B/-^*livers following combinatorial treatment may be a sign of improved liver development following rescue of liver pathology by *SMN1* restoration ^64,65^. Additionally, mifepristone has been shown to reduce liver enzymes in patients with non-alcoholic fatty liver disease and could be beneficial knowing that scAAV9-*SMN1* can cause liver toxicity ^45^.

Interestingly, we found an increase in myofiber area only in *Smn^2B/-^*females following combinatorial treatment while there were no changes in males. This may be due to previously reported sex-dependent differences in both muscle metabolism and GC-GR activity ^66–68^. Fundamentally, the beneficial effects of mifepristone have been associated with skeletal muscle pathology. We demonstrate that combinatorial therapy can improve certain aspects of disease pathology beyond that of treatment with the gene therapy alone in *Smn^2B/-^* mice. As a result of its success so far, scAAV9-*SMN1* now requires long-term assessment as it is possible that patients living longer may experience muscle and metabolic pathologies that need to be addressed therapeutically ^69^.

Metabolic pathologies in SMA have been reported, all be it to a limited extent, since before the genetic discovery of the *SMN* gene ^4^. A more recent nutritional study found that nusinersen-treated patients still have gastrointestinal issues that can be modulated by an amino acid diet ^18^. Additionally, therapies targeting complementary pathways that ultimately increase SMN in peripheral tissues prolong survival, demonstrating the importance of peripheral rescue beyond the canonical pathologies of SMA ^70^. Along with the fact that the three approved gene therapies for SMA are unfortunately not cures for the disease, it is now clear that metabolic defects are often motor neuron-independent and should be autonomously addressed ^71^.

Notably, there are many commonalities in metabolic pathologies between Cushing’s syndrome (CS), a serious endocrine disorder, and SMA, including hyperglycaemia, fatty liver and muscle atrophy. FDA-approved mifepristone (Corlux^®^) is currently a widely used therapy for CS. In addition, both respiratory and digestive system dysfunctions have been associated with the prevalence of depression in SMA patients ^72^. Psychological disorders including anxiety and depression often accompany chronic disease and mifepristone has been investigated for its therapeutic benefit in patients with psychotic depression. Mifepristone-treated patients had reduced psychotic symptoms compared to placebo-treated patients, with a large safety margin ^73^. Consequently, mifepristone could address metabolic pathologies, mental health issues, and neural SMN independent pathologies, in SMA patients through its ability to cross the blood-brain barrier ^74^.

In summary, our results along with our previous work ^3^, suggest that the GC-GR-*Klf15* pathway is dysregulated in metabolic tissues of SMA patients and mouse models and could play a role in glucose, lipid and amino acid metabolic dysfunctions ^3^. Following ground-breaking research and clinical trials, the approval of three gene-directed therapies has revolutionised the field of SMA. With that said, it is now time to utilise these therapies to maximise potential therapeutic benefit. Additional peripheral and metabolic pathologies are important targets for therapeutic interventions that have until recently received little focus ^4^. Future investigations should therefore be aimed at furthering our understanding of SMN-independent contributors to SMA pathology.

## DATA AVAILABILITY STATEMENT

All data associated with this study are available in the main text or supplementary materials. Raw data can be provided upon request.

## ACKNOWLEDGEMENTS

The authors would like to thank the staff at the Biomedical Sciences Unit (BSU) facility at the University of Keele as well as Biological Research Resources (BRR) staff at the University of Edinburgh for excellent animal care and husbandry

## AUTHOR CONTRIBUTIONS

Conceptualization: E.R.S, M.B; Methodology: E.R.S, P.P.T, M.D, S.D, B.L.S & M.B; Formal Analysis: E.R.S, P.P.T, M.D, H.C, Y.T.H & M.B; Investigation: E.R.S, E.M, J.M.H, O.C, P.P.T, M.D, H.C, Y.T.H, T.H.G, S.B, L.C, T.S, & M.B; Software: E.R.S; Visualisation: E.R.S; Resources: M,D, T.H.G & M.B; Writing – Original Draft: E.R.S, M.B; writing – Review and Editing: E.R.S, E.M, J.M.H, O.C, P.P.T, M.D, H.C, Y.T.H, T.H.G, S.B, L.C, T.S, S.D, B.L.S & M.B ; Supervision: M.B; Project Administration: M.B; Funding Acquisition: M.B.

## REFERENCES

1 Lefebvre S, Bürglen L, Reboullet S, Clermont O, Burlet P, Viollet L et al. Identification and characterization of a spinal muscular atrophy-determining gene. Cell 1995; 80: 155– 165.

2 Hua Y, Sahashi K, Rigo F, Hung G, Horev G, Bennett CF et al. Peripheral SMN restoration is essential for long-term rescue of a severe spinal muscular atrophy mouse model. Nature 2011; 478: 123–126.

3 Walter LM, Deguise M-O, Meijboom KE, Betts CA, Ahlskog N, van Westering TLE et al. Interventions Targeting Glucocorticoid-Krüppel-like Factor 15-Branched-Chain Amino Acid Signaling Improve Disease Phenotypes in Spinal Muscular Atrophy Mice. EBioMedicine 2018; 31: 226–242.

4 Deguise M-O, Chehade L, Kothary R. Metabolic Dysfunction in Spinal Muscular Atrophy. Int J Mol Sci 2021; 22: 5913.

5 Pearn J. Classification of spinal muscular atrophies. Lancet Lond Engl 1980; 1: 919– 922.

6 Crawford TO, Pardo CA. The neurobiology of childhood spinal muscular atrophy. Neurobiol Dis 1996; 3: 97–110.

7 Lorson CL, Hahnen E, Androphy EJ, Wirth B. A single nucleotide in the SMN gene regulates splicing and is responsible for spinal muscular atrophy. Proc Natl Acad Sci U S A 1999; 96: 6307–6311.

8 d’Ydewalle C, Sumner CJ. Spinal Muscular Atrophy Therapeutics: Where do we Stand? Neurotherapeutics 2015; 12: 303–316.

9 Bowerman M, Becker CG, Yáñez-Muñoz RJ, Ning K, Wood MJA, Gillingwater TH et al. Therapeutic strategies for spinal muscular atrophy: SMN and beyond. Dis Model Mech 2017; 10: 943–954.

10 Deguise M-O, Baranello G, Mastella C, Beauvais A, Michaud J, Leone A et al. Abnormal fatty acid metabolism is a core component of spinal muscular atrophy. Ann Clin Transl Neurol 2019; 6: 1519–1532.

11 Yeo CJJ, Darras BT. Overturning the Paradigm of Spinal Muscular Atrophy as Just a Motor Neuron Disease. Pediatr Neurol 2020; 109: 12–19.

12 Yeo CJJ, Levine A, Darras B. Hepatic Steatosis in Patients with Spinal Muscular Atrophy (SMA) (P1-1.Virtual). Neurology 2022; 98.https://n.neurology.org/content/98/18_Supplement/2692 (accessed 17 Jul2022).

13 Bowerman M, Swoboda KJ, Michalski J-P, Wang G-S, Reeks C, Beauvais A et al. Glucose Metabolism and Pancreatic Defects in Spinal Muscular Atrophy. Ann Neurol 2012; 72: 256–268.

14 Djordjevic SA, Milic-Rasic V, Brankovic V, Kosac A, Dejanovic-Djordjevic I, Markovic-Denic L et al. Glucose and lipid metabolism disorders in children and adolescents with spinal muscular atrophy types 2 and 3. Neuromuscul Disord NMD 2021; 31: 291–299.

15 Narver HL, Kong L, Burnett BG, Choe DW, Bosch-Marcé M, Taye AA et al. Sustained improvement of spinal muscular atrophy mice treated with trichostatin a plus nutrition. Ann Neurol 2008; 64: 465–470.

16 Butchbach MER, Rose FF, Rhoades S, Marston J, Mccrone JT, Sinnott R et al. Effect of diet on the survival and phenotype of a mouse model for spinal muscular atrophy. Biochem Biophys Res Commun Biochem Biophys Res Commun January 2010; 1: 835– 840.

17 Butchbach MER, Singh J, Gurney ME, Burghes AHM. The effect of diet on the protective action of D156844 observed in spinal muscular atrophy mice. Exp Neurol 2014; 256: 1– 6.

18 O’Connor G, Edel L, Raquq S, Bowerman M, Szmurlo A, Simpson Z et al. Open-labelled study to monitor the effect of an amino acid formula on symptom management in children with spinal muscular atrophy type I: The SMAAF pilot study. Nutr Clin Pract 2023; 38: 871–880.

19 Sutton ER, Beauvais A, Yaworski R, De Repentigny Y, Reilly A, Alves de Almeida MM et al. Liver SMN restoration rescues the Smn2B/-mouse model of spinal muscular atrophy. EBioMedicine 2024; 110: 105444.

20 Groen EJN, Talbot K, Gillingwater TH. Advances in therapy for spinal muscular atrophy: promises and challenges. Nat Rev Neurol 2018; 14: 214–224.

21 Wood MJA, Talbot K, Bowerman M. Spinal muscular atrophy: antisense oligonucleotide therapy opens the door to an integrated therapeutic landscape. Hum Mol Genet 2017; 26: R151–R159.

22 Gray S, Feinberg MW, Hull S, Kuo CT, Watanabe M, Sen-Banerjee S et al. The Krüppel-like factor KLF15 regulates the insulin-sensitive glucose transporter GLUT4. J Biol Chem 2002; 277: 34322–34328.

23 Gray S, Wang B, Orihuela Y, Hong E-G, Fisch S, Haldar S et al. Regulation of gluconeogenesis by Krüppel-like factor 15. Cell Metab 2007; 5: 305–312.

24 Jeyaraj D, Scheer FAJL, Ripperger JA, Haldar SM, Lu Y, Prosdocimo DA et al. Klf15 orchestrates circadian nitrogen homeostasis. Cell Metab 2012; 15: 311–323.

25 Haldar SM, Jeyaraj D, Anand P, Zhu H, Lu Y, Prosdocimo DA et al. Kruppel-like factor 15 regulates skeletal muscle lipid flux and exercise adaptation. Proc Natl Acad Sci U S A 2012; 109: 6739–6744.

26 Masuno K, Haldar SM, Jeyaraj D, Mailloux CM, Huang X, Panettieri RA et al. Expression profiling identifies Klf15 as a glucocorticoid target that regulates airway hyperresponsiveness. Am J Respir Cell Mol Biol 2011; 45: 642–9.

27 Shimizu N, Yoshikawa N, Ito N, Maruyama T, Suzuki Y, Takeda S et al. Crosstalk between Glucocorticoid Receptor and Nutritional Sensor mTOR in Skeletal Muscle. Cell Metab 2011; 13: 170–182.

28 Meijboom KE, Volpato V, Monzón-Sandoval J, Hoolachan JM, Hammond SM, Abendroth F et al. Combining multiomics and drug perturbation profiles to identify muscle-specific treatments for spinal muscular atrophy. 2021. doi:10.1172/jci.insight.149446.

29 Mathew S, Ticsa MS, Qadir S, Rezene A, Khanna D. Multiple Clinical Indications of Mifepristone: A Systematic Review. Cureus 2023; 15: e48372.

30. 30 005058 - SMA-like Strain Details. https://www.jax.org/strain/005058 (accessed 12 Aug2022).

31 Gaudry J-P, Aebi A, Valdés P, Schneider BL. Production and Purification of Adeno-Associated Viral Vectors (AAVs) Using Orbitally Shaken HEK293 Cells. In: Hacker DL (ed). Recombinant Protein Expression in Mammalian Cells: Methods and Protocols. Springer US: New York, NY, 2024, pp 55–74.

32 Briese M, Esmaeili B, Fraboulet S, Burt EC, Christodoulou S, Towers PR et al. Deletion of smn-1, the Caenorhabditis elegans ortholog of the spinal muscular atrophy gene, results in locomotor dysfunction and reduced lifespan. Hum Mol Genet 2009; 18: 97– 104.

33 Dimitriadi M, Derdowski A, Kalloo G, Maginnis MS, O’Hern P, Bliska B, et al. Decreased function of survival motor neuron protein impairs endocytic pathways. Proc Natl Acad Sci 2016; 113: E4377–E4386.

34 Yaffe D, Saxel O. Serial passaging and differentiation of myogenic cells isolated from dystrophic mouse muscle. Nature 1977; 270: 725–727.

35 Green H, Meuth M. An established pre-adipose cell line and its differentiation in culture. Cell 1974; 3: 127–133.

36 Breslow JL, Sloan HR, Ferrans VJ, Anderson JL, Levy RI. Characterization of the mouse liver cell line FL83B. Exp Cell Res 1973; 78: 441–453.

37 Hinds TD, Ramakrishnan S, Cash HA, Stechschulte LA, Heinrich G, Najjar SM et al. Discovery of glucocorticoid receptor-beta in mice with a role in metabolism. Mol Endocrinol Baltim Md 2010; 24: 1715–1727.

38 Hinds TD, Peck B, Shek E, Stroup S, Hinson J, Arthur S et al. Overexpression of Glucocorticoid Receptor β Enhances Myogenesis and Reduces Catabolic Gene Expression. Int J Mol Sci 2016; 17. doi:10.3390/ijms17020232.

39 Bowerman M, Murray LM, Beauvais A, Pinheiro B, Kothary R. A critical smn threshold in mice dictates onset of an intermediate spinal muscular atrophy phenotype associated with a distinct neuromuscular junction pathology. Neuromuscul Disord NMD 2012; 22: 263–276.

40 Hsieh-Li HM, Chang JG, Jong YJ, Wu MH, Wang NM, Tsai CH et al. A mouse model for spinal muscular atrophy. Nat Genet 2000; 24: 66–70.

41 Ahlskog N, Hayler D, Krueger A, Kubinski S, Claus P, Hammond SM et al. Muscle overexpression of Klf15 via an AAV8-Spc5-12 construct does not provide benefits in spinal muscular atrophy mice. Gene Ther 2020; 27: 505–515.

42 Dimitriadi M, Kye MJ, Kalloo G, Yersak JM, Sahin M, Hart AC. The neuroprotective drug riluzole acts via small conductance Ca2+-activated K+ channels to ameliorate defects in spinal muscular atrophy models. J Neurosci Off J Soc Neurosci 2013; 33: 6557–6562.

43 Hoolachan JM, McCallion E, Sutton ER, Çetin Ö, Pacheco-Torres P, Dimitriadi M et al. A transcriptomics-based drug repositioning approach to identify drugs with similar activities for the treatment of muscle pathologies in spinal muscular atrophy (SMA) models. Hum Mol Genet 2023; : ddad192.

44 Roussos A, Kitopoulou K, Borbolis F, Palikaras K. Caenorhabditis elegans as a Model System to Study Human Neurodegenerative Disorders. Biomolecules 2023; 13: 478.

45 Ragucci E, Nguyen D, Lamerson M, Moraitis AG. Effects of Mifepristone on Nonalcoholic Fatty Liver Disease in a Patient with a Cortisol-Secreting Adrenal Adenoma. Case Rep Endocrinol 2017; 2017: e6161348.

46 Reilly A, Deguise M-O, Beauvais A, Yaworski R, Thebault S, Tessier DR et al. Central and peripheral delivered AAV9-SMN are both efficient but target different pathomechanisms in a mouse model of spinal muscular atrophy. Gene Ther 2022. doi:10.1038/s41434-022-00338-1.

47 Bose SK, Hutson I, Harris CA. Hepatic Glucocorticoid Receptor Plays a Greater Role Than Adipose GR in Metabolic Syndrome Despite Renal Compensation. Endocrinology 2016; 157: 4943–4960.

48 Zinker B, Mika A, Nguyen P, Wilcox D, Öhman L, Geldern TW von et al. Liver-selective glucocorticoid receptor antagonism decreases glucose production and increases glucose disposal, ameliorating insulin resistance. Metab - Clin Exp 2007; 56: 380–387.

49 Nabatame Y, Hosooka T, Aoki C, Hosokawa Y, Imamori M, Tamori Y et al. Kruppel-like factor 15 regulates fuel switching between glucose and fatty acids in brown adipocytes. J Diabetes Investig 2021; 12: 1144–1151.

50 Nagare T, Sakaue H, Matsumoto M, Cao Y, Inagaki K, Sakai M et al. Overexpression of KLF15 Transcription Factor in Adipocytes of Mice Results in Down-regulation of SCD1 Protein Expression in Adipocytes and Consequent Enhancement of Glucose-induced Insulin Secretion*. J Biol Chem 2011; 286: 37458–37469.

51 Gubbi S, Muniyappa R, Sharma ST, Grewal S, McGlotten R, Nieman LK. Mifepristone Improves Adipose Tissue Insulin Sensitivity in Insulin Resistant Individuals. J Clin Endocrinol Metab 2021; 106: 1501–1515.

52 Díaz-Castro F, Monsalves-Álvarez M, Rojo LE, Campo A del, Troncoso R. Mifepristone for Treatment of Metabolic Syndrome: Beyond Cushing’s Syndrome. Front Pharmacol 2020; 11. doi:10.3389/fphar.2020.00429.

53 Britto FA, Begue G, Rossano B, Docquier A, Vernus B, Sar C et al. REDD1 deletion prevents dexamethasone-induced skeletal muscle atrophy. Am J Physiol Endocrinol Metab 2014; 307: E983–993.

54 Gordon BS, Liu C, Steiner JL, Nader GA, Jefferson LS, Kimball SR. Loss of REDD1 augments the rate of the overload-induced increase in muscle mass. Am J Physiol - Regul Integr Comp Physiol 2016; 311: R545–R557.

55 Bowerman M, Beauvais A, Anderson CL, Kothary R. Rho-kinase inactivation prolongs survival of an intermediate SMA mouse model. Hum Mol Genet 2010; 19: 1468–1478.

56 Dangouloff T, Servais L. Clinical Evidence Supporting Early Treatment Of Patients With Spinal Muscular Atrophy: Current Perspectives. Ther Clin Risk Manag 2019; 15: 1153– 1161.

57 Shih ST, Farrar MA, Wiley V, Chambers G. Newborn screening for spinal muscular atrophy with disease-modifying therapies: a cost-effectiveness analysis. J Neurol Neurosurg Psychiatry 2021; 92: 1296–1304.

58 Dominguez E, Marais T, Chatauret N, Benkhelifa-Ziyyat S, Duque S, Ravassard P et al. Intravenous scAAV9 delivery of a codon-optimized SMN1 sequence rescues SMA mice. Hum Mol Genet 2011; 20: 681–693.

59 Reilly A, Yaworski R, Beauvais A, Schneider BL, Kothary R. Long term peripheral AAV9-SMN gene therapy promotes survival in a mouse model of spinal muscular atrophy. Hum Mol Genet 2023; : ddad202.

60 Reilly A, Chehade L, Kothary R. Curing SMA: Are we there yet? Gene Ther 2023; 30: 8– 17.

61 Marrugal-Lorenzo JA, Serna-Gallego A, González-González L, Buñuales M, Poutou J, Pachón J et al. Inhibition of adenovirus infection by mifepristone. Antiviral Res 2018; 159: 77–83.

62 Cuevas-Ramos D, Lim DST, Fleseriu M. Update on medical treatment for Cushing’s disease. Clin Diabetes Endocrinol 2016; 2: 16.

63 Grunberg SM, Weiss MH, Russell CA, Spitz IM, Ahmadi J, Sadun A et al. Long-term administration of mifepristone (RU486): clinical tolerance during extended treatment of meningioma. Cancer Invest 2006; 24: 727–733.

64 Szunyogova E, Zhou H, Maxwell GK, Powis RA, Francesco M, Gillingwater TH et al. Survival Motor Neuron (SMN) protein is required for normal mouse liver development. Sci Rep 2016; 6. doi:10.1038/srep34635.

65 Anzai K, Tsuruya K, Ida K, Kagawa T, Inagaki Y, Kamiya A. Kruppel-like factor 15 induces the development of mature hepatocyte-like cells from hepatoblasts. Sci Rep 2021; 11: 18551.

66 Rosa-Caldwell ME, Greene NP. Muscle metabolism and atrophy: let’s talk about sex. Biol Sex Differ 2019; 10: 43.

67 Li S, Schönke M, Buurstede JC, Moll TJA, Gentenaar M, Schilperoort M et al. Sexual Dimorphism in Transcriptional and Functional Glucocorticoid Effects on Mouse Skeletal Muscle. Front Endocrinol 2022; 13: 907908.

68 Yoshikawa N, Oda A, Yamazaki H, Yamamoto M, Kuribara-Souta A, Uehara M et al. The Influence of Glucocorticoid Receptor on Sex Differences of Gene Expression Profile in Skeletal Muscle. Endocr Res 2021; 46: 99–113.

69 Stevens D, Claborn MK, Gildon BL, Kessler TL, Walker C. Onasemnogene Abeparvovec-xioi: Gene Therapy for Spinal Muscular Atrophy. Ann Pharmacother 2020; 54: 1001–1009.

70 Dumas SA, Villalón E, Bergman EM, Wilson KJ, Marugan JJ, Lorson CL et al. A combinatorial approach increases SMN level in SMA model mice. Hum Mol Genet 2022; : ddac068.

71 Bryda EC. The Mighty Mouse: The Impact of Rodents on Advances in Biomedical Research. Mo Med 2013; 110: 207–211.

72 Yao M, Xia Y, Feng Y, Ma Y, Hong Y, Zhang Y et al. Anxiety and depression in school-age patients with spinal muscular atrophy: a cross-sectional study. Orphanet J Rare Dis 2021; 16: 385.

73 Block TS, Kushner H, Kalin N, Nelson C, Belanoff J, Schatzberg A. Combined Analysis of Mifepristone for Psychotic Depression: Plasma Levels Associated With Clinical Response. Biol Psychiatry 2018; 84: 46–54.

74 Check JH, Wilson C, Cohen R, Sarumi M. Evidence that Mifepristone, a Progesterone Receptor Antagonist, Can Cross the Blood Brain Barrier and Provide Palliative Benefits for Glioblastoma Multiforme Grade IV. Anticancer Res 2014; 34: 2385–2388.

