## Supplementary figures and images for "Mifepristone alone and in combination with scAAV9-*SMN1* gene therapy improves disease phenotypes in *Smn^2B/-^* spinal muscular atrophy mice"

### Supplementary Figure 1

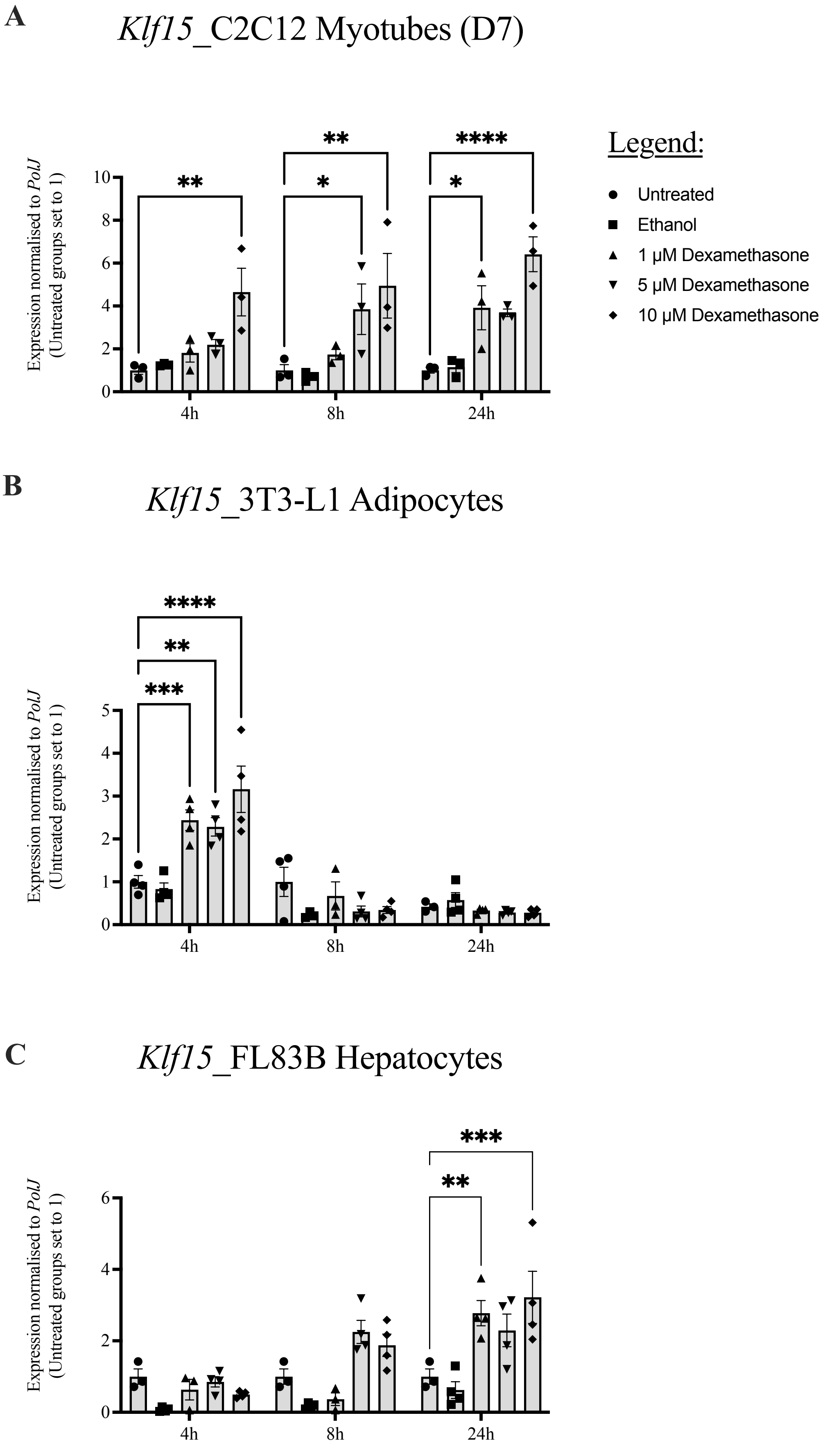

### Supplementary Figure 2

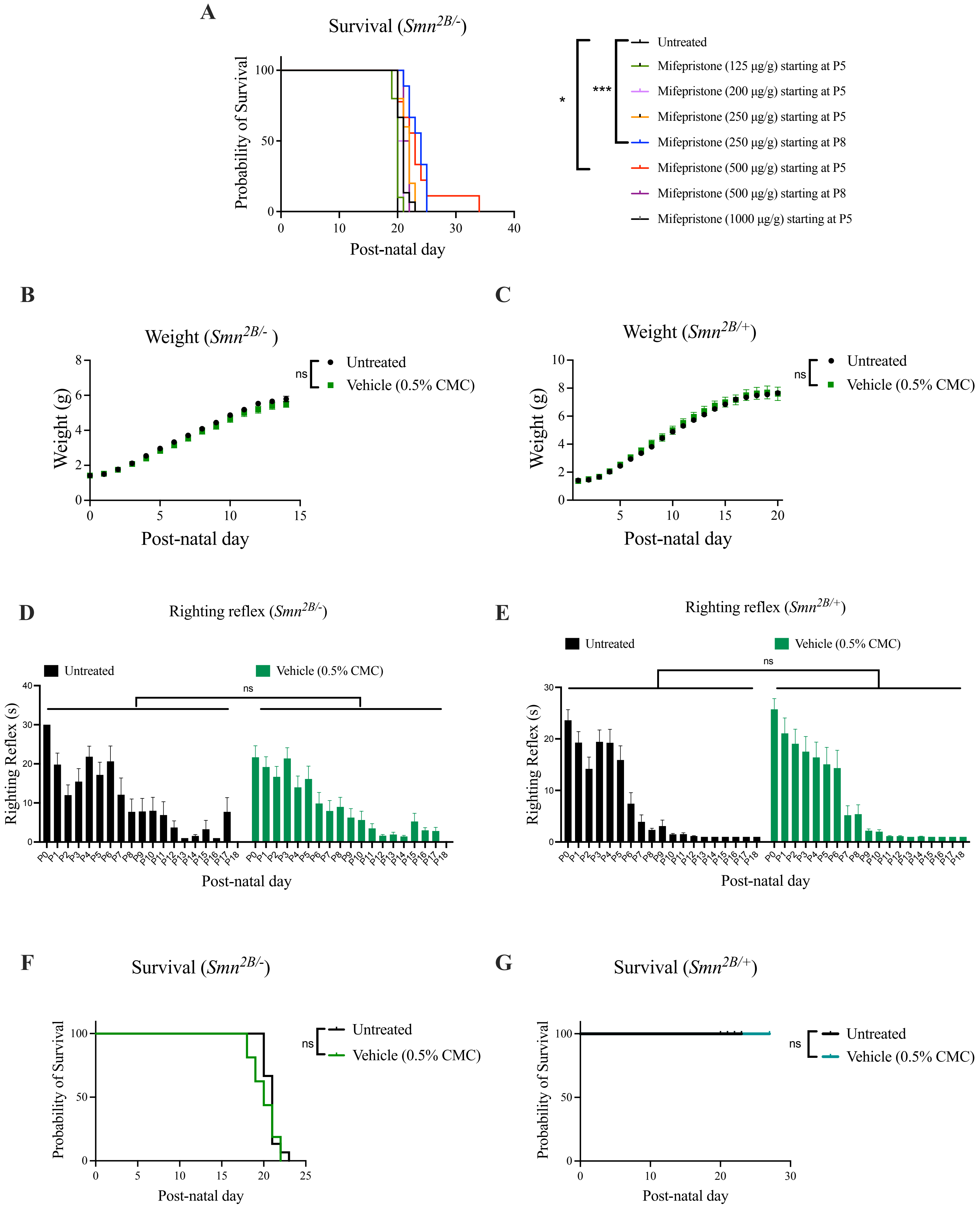

### Supplementary Figure 3

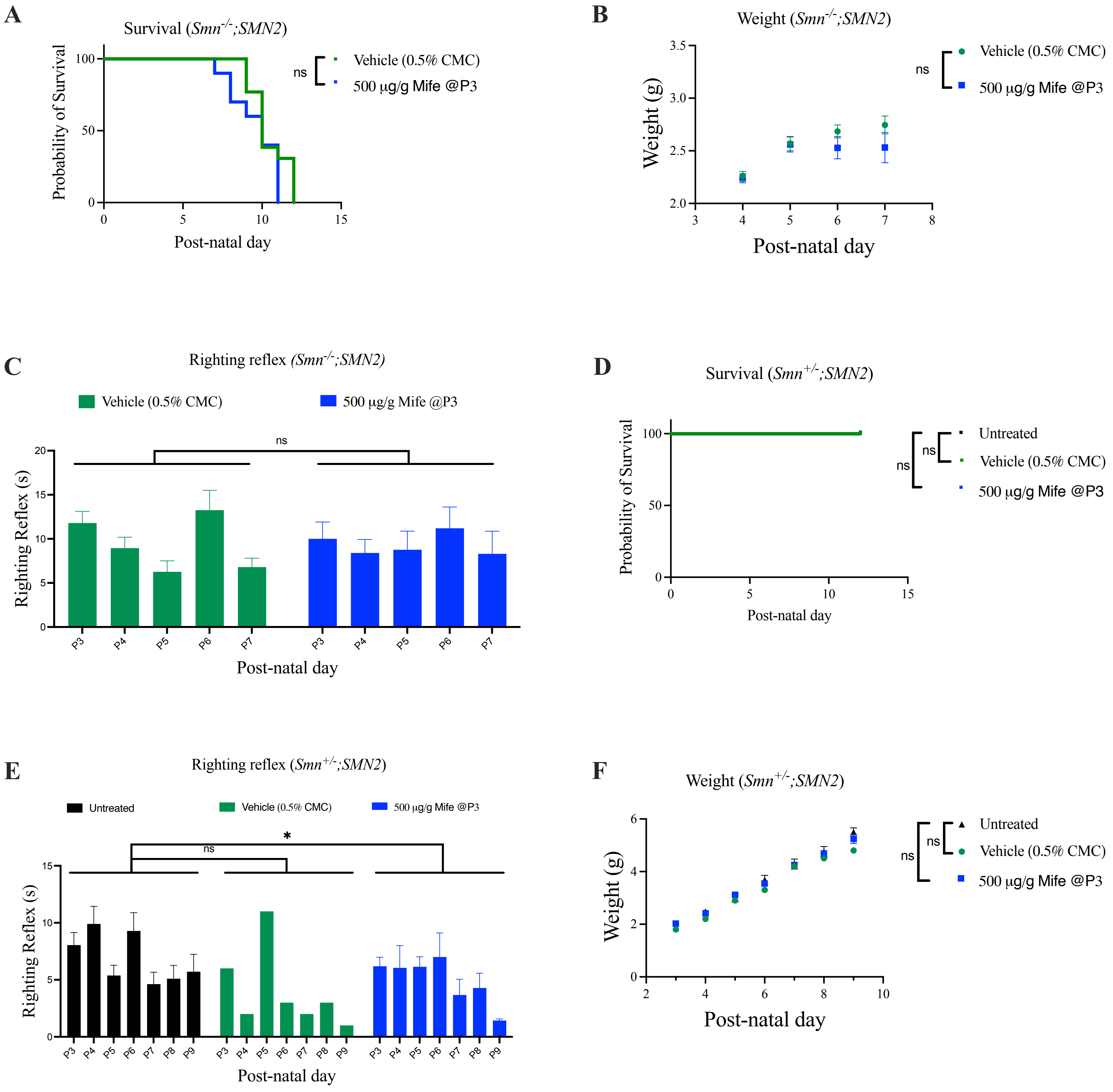

### Supplementary Figure 4

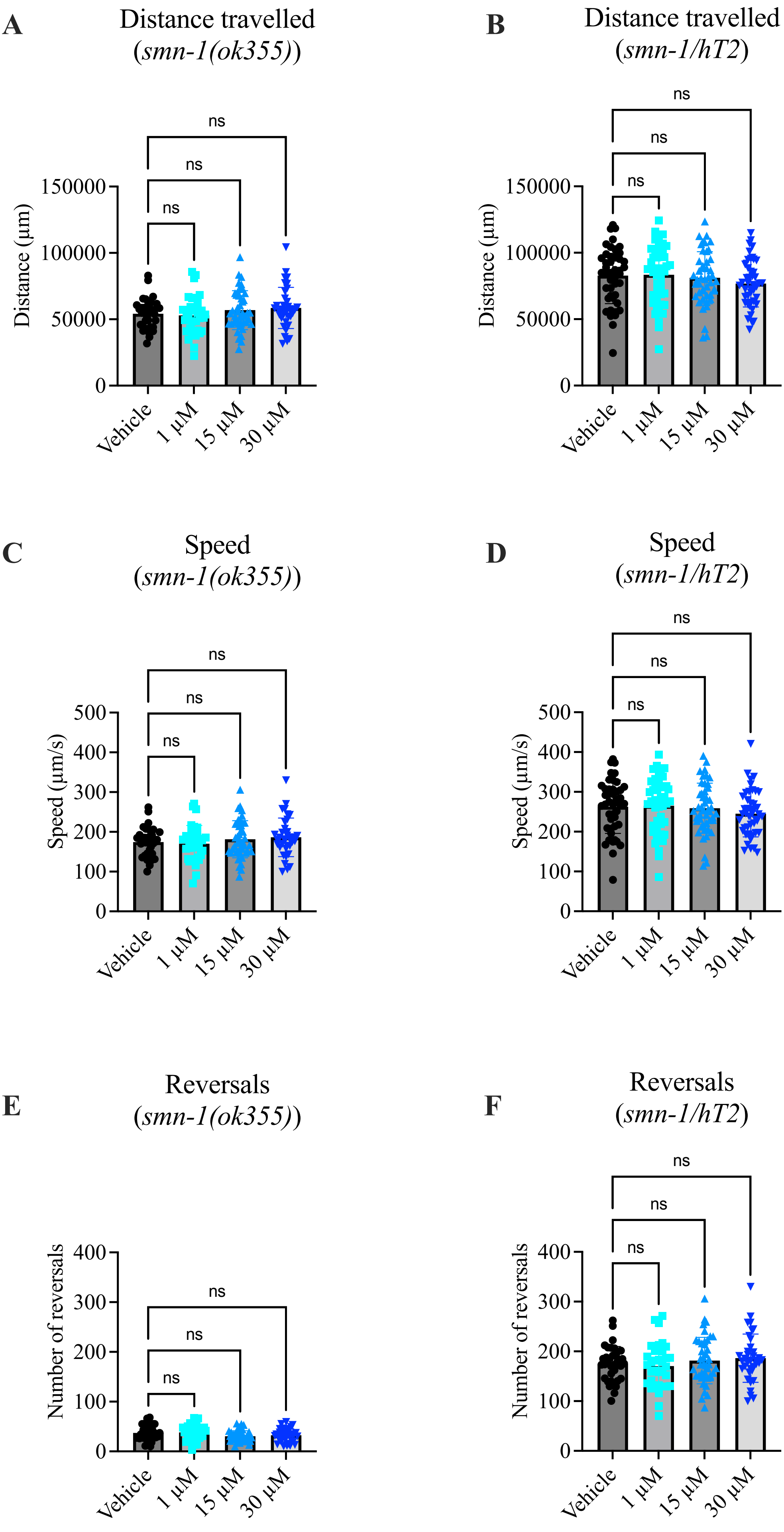

### Supplementary Figure 5

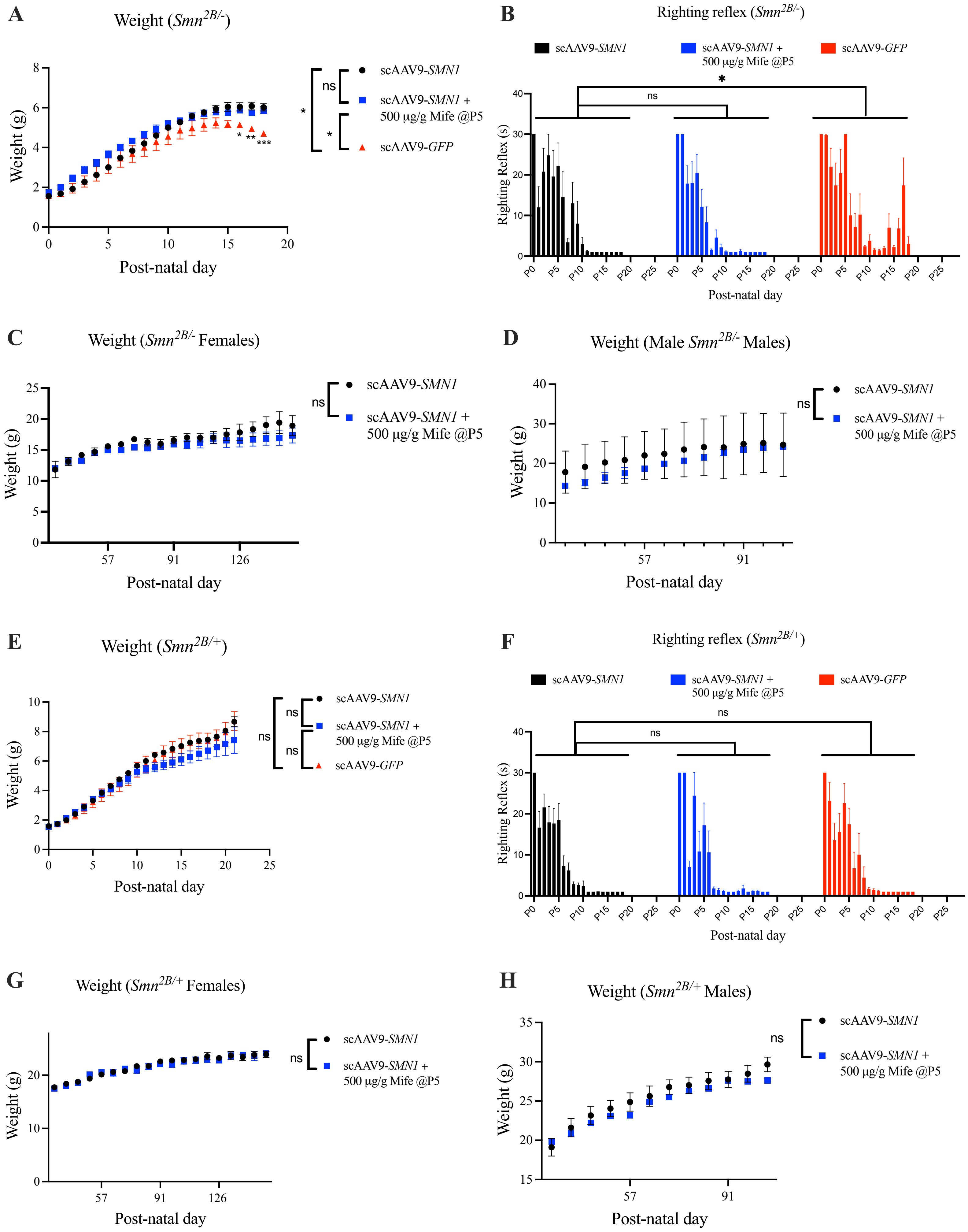
