## Supplementary Table 1 for "Mifepristone alone and in combination with scAAV9-*SMN1* gene therapy improves disease phenotypes in *Smn^2B/-^* spinal muscular atrophy mice"

**Supplementary Table 1. List of qPCR primers used.**

| **Gene** | **Forward primer** | **Reverse primer** |
| --- | --- | --- |
| ***PolJ (mouse)*** | ACCACACTCTGGGGAACA TC | CTCGCTGA TGAGGTCTGTGA |
| ***GRα (mouse)*** | AAAGAGCTAGGAAAAGCCATTGTC | TCAGCTAACATCTCTGGGAATTCA |
| ***GRβ (mouse)*** | AAAGAGCTAGGAAAAGCCATTGTC | CTGTCTTTGGGCTTTTGAGATAGG |
| ***Klf15 (mouse)*** | TGCGTCGGCACACAGGCGAGAA | CCGGTGCCTTGACAACTCA TCT |
| ***Pol2RA (human)*** | CAACGCACACATCCAGAACG | TCCTTGACTCCCTCCACCAC |
| ***Klf15 (human)*** | GCTTGAGTTAAATGTGCAGGG | TTCTAAATCAGGGTTGGGAGG |
